# Binding kinetics enable tumor-selective MAPK and growth suppression by a Type 1 RAF Inhibitor

**DOI:** 10.64898/2026.10.05.756774

**Authors:** Mathieu Desaunay, Tara L. Peters, Evangelia Matenoglou, Beau Baars, Bijaya Gaire, Ana Orive-Ramos, Li Ren, Joseph P. Lyssikatos, Michael R. Burkard, Dalton Dacus, Matthew J. Sale, Stuart A. Aaronson, Frank McCormick, Evripidis Gavathiotis, Stefan D. Gross, Poulikos I. Poulikakos

## Abstract

Therapeutic inhibition of the RAS/MAPK cascade, an oncogenic driver in over one-third of human cancers, has been limited by a core trade-off: effective pathway blockade in tumors versus dose-limiting toxicity in normal tissues. Although Type 1.5 RAF inhibitors successfully mitigated this trade-off in BRAF-monomer–driven tumors, an equivalent strategy has been lacking for RAF-dimer– driven cancers, including those with RAS mutations. Here, we characterize ELV-3111, a next-generation, potent, and selective Type 1 RAF inhibitor that overcomes this limitation via a distinct mechanism of action. ELV-3111 induces paradoxical MAPK hyperactivation in normal tissues by binding RAF and promoting the active conformation (priming) followed by rapid dissociation (fast off-rate), a biochemical feature that spares RAS-mutant tumors, where RAF is already primed. In these and other RAF dimer-driven tumor models, ELV-3111 achieves potent and selective MAPK and growth suppression. When combined with a MEK inhibitor, ELV-3111 enables complementary pharmacology: additive inhibition in tumors and compensatory effects in normal tissues. The combination produced durable regressions across RAS-mutant and BRAF-mutant models, including a RAS-mutant model refractory to current therapies, while maintaining favorable tolerability. Thus, we uncover a generalizable mechanism that enables Type 1 RAF inhibitor plus MEK inhibitor combinations to achieve tumor-selective MAPK suppression, with the potential to inform targeted therapy design.

## Introduction

Over a third of human tumors are driven by deregulated RAS/MAPK signaling, most commonly associated with mutations in RAS or BRAF(*1–3*). RAS/MAPK-targeting therapies, including direct RAS inhibitors and small-molecule inhibitors of RAF, MEK and ERK, have been developed as effective treatments for these cancers. Direct mutant-selective KRAS inhibitors represent a major recent advance and have demonstrated meaningful clinical benefit with generally favorable tolerability. However, adaptive and acquired resistance frequently restore MAPK signaling, resulting in disease progression in most patients. More broadly, despite clinical activity, the efficacy of RAS/MAPK-targeting therapies is often limited by a therapeutic ceiling, whereby toxicities emerge at doses that remain insufficient to achieve complete and durable MAPK pathway suppression in tumors.

Protein kinase inhibitors are structurally classified according to the conformation of the DFG motif and αC-helix that they stabilize, including Type 1 (DFG-IN, αC-IN), Type 1.5 (DFG-IN, αC-OUT), and Type 2 (DFG-OUT, αC-IN) inhibitors(*4*). In RAF kinases, these structural classes have distinct functional consequences. We previously showed that the position of the αC-helix determines inhibitor selectivity toward RAF monomers versus dimers: αC-OUT Type 1.5 inhibitors selectively inhibit RAF monomers, whereas αC-IN Type 1 and Type 2 inhibitors inhibit both RAF monomers and dimers(*5, 6*).

Current clinically approved RAF inhibitors belong to the Type 1.5 class and selectively inhibit monomeric BRAF(V600X)(*7*). In cells with dimeric RAF, including normal cells, these RAF inhibitors paradoxically activate RAF and MAPK signaling via an allosteric mechanism described previously by us(*5, 8, 9*) and others(*10*). This opposing pharmacology (MAPK inhibition in tumors and paradoxical MAPK activation in normal tissues) enabled combinations with MEK inhibitors (MEKis)(*11–13*) or EGFR antagonists(*14*), enhancing efficacy and tolerability by simultaneously suppressing MAPK signaling in tumors while buffering pathway inhibition in normal tissues. Consequently, combined RAF and MEK inhibition has become the standard of care for BRAF(V600X) tumors.

Despite their clinical success, their inactivity in dimeric RAF contexts restricted the therapeutic utility of Type 1.5 RAF inhibitors to BRAF(V600X) tumors. To broaden therapeutic utility, Type 2 RAF inhibitors, which inhibit both monomeric and dimeric RAF, were developed to target a wider spectrum of RAS/MAPK-driven tumors, including RAS-mutant cancers. However, as single agents, Type 2 RAFis have shown limited clinical activity(*15, 16*), necessitating combinations with MEK inhibitors. In contrast to Type 1.5 RAF inhibitors, however, both agents suppress MAPK signaling in normal tissues, narrowing the therapeutic index. Consequently, dose reductions required to manage toxicity compromise pathway inhibition in tumors, yielding predominantly stable disease rather than durable tumor regressions(*17*). Type 1 RAF inhibitors, which also inhibit both monomeric and dimeric RAF, were among the earliest RAF inhibitors developed, but none advanced beyond early clinical development because of insufficient potency and selectivity for therapeutic use.

Here, we show that ELV-3111, a novel, potent and selective Type 1 RAF inhibitor, effectively overcomes key limitations of both Type 1.5 and Type 2 RAF inhibitors. ELV-3111 potently inhibits BRAF class I/II/III-, CRAF-, and RAS-mutant cancer models *in vitro* and *in vivo* and overcomes multiple mechanisms of resistance to current RAF inhibitors, including BRAF(V600E) splice variants and RAS co-mutations. Unlike Type 2 RAF inhibitors, ELV-3111 induces robust paradoxical MAPK hyperactivation in normal cells through a distinct mechanism involving RAS-dependent RAF priming upon inhibitor binding followed by rapid inhibitor dissociation. This biochemical and pharmacologic profile forms the basis for a therapeutic strategy combining a Type 1 RAF inhibitor with MEK inhibition that produces robust tumor control in RAS- and BRAF-mutant models, including RAS-mutant tumors refractory to current MAPK-targeting therapies.

Thus, we uncover a previously unrecognized mechanism of paradoxical MAPK activation by Type 1 RAF inhibitors that integrates the therapeutic advantages of both Type 1.5 and Type 2 RAF inhibitors, extending the therapeutic principle underlying RAF/MEK inhibition in BRAF(V600X) tumors to RAF dimer-driven cancers, including RAS-mutant tumors.

## Results

### ELV-3111 is a Type 1 RAF inhibitor with broad activity and potent inhibition of both monomeric and dimeric RAF

ELV-3111 (**Fig. 1A and fig. S1, A to F**) was discovered and developed through optimization of a chemical series of BRAF inhibitors, and is a potent, highly selective BRAF inhibitor based on a scaffold that binds the kinase active site in a Type 1 binding (*4*). A crystal structure of ELV-3111 and the BRAF kinase domain solved to a 2.85 Å resolution (**fig. S1, G and H and Table S1)** revealed that ELV-3111 binds BRAF in a DFG-IN/αC-helix-IN (Type 1) binding conformation (**Fig. 1B**). To understand the selectivity of ELV-3111 versus the broader kinome, the compound was profiled in biochemical assays against a panel of 372 protein kinases at a concentration of 1μM. Only 17 of these kinases were inhibited by more than 50% and were therefore considered potential anti-targets (**Fig. 1C and Table S2**). The IC_50_ values of ELV-3111 for each of these 17 kinases were determined, and ELV-3111 was found to be 17 to 844-fold more selective for BRAF over any of the potential off-target kinases (**Table S2**). Some off-targets identified in the kinase screen, such as ABL1 or c-KIT, are known to cause toxicity in humans, so for a subset of these, we also determined cellular IC_50_s in relevant functional assays. Results demonstrated that in cells, ELV-3111 is at least 40-fold more selective for BRAF than these anti-targets (**Table S2**).

**Fig. 1.**
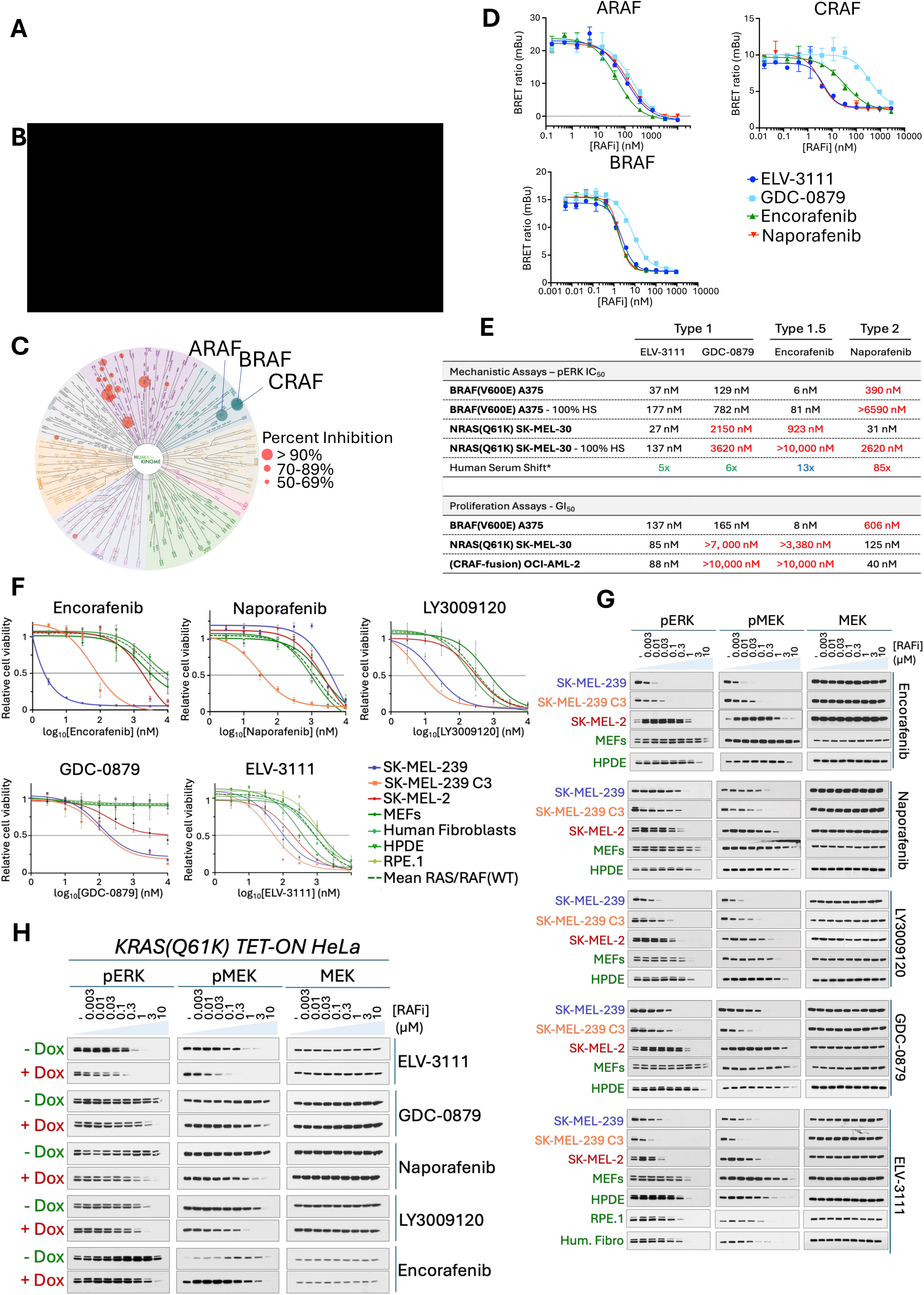
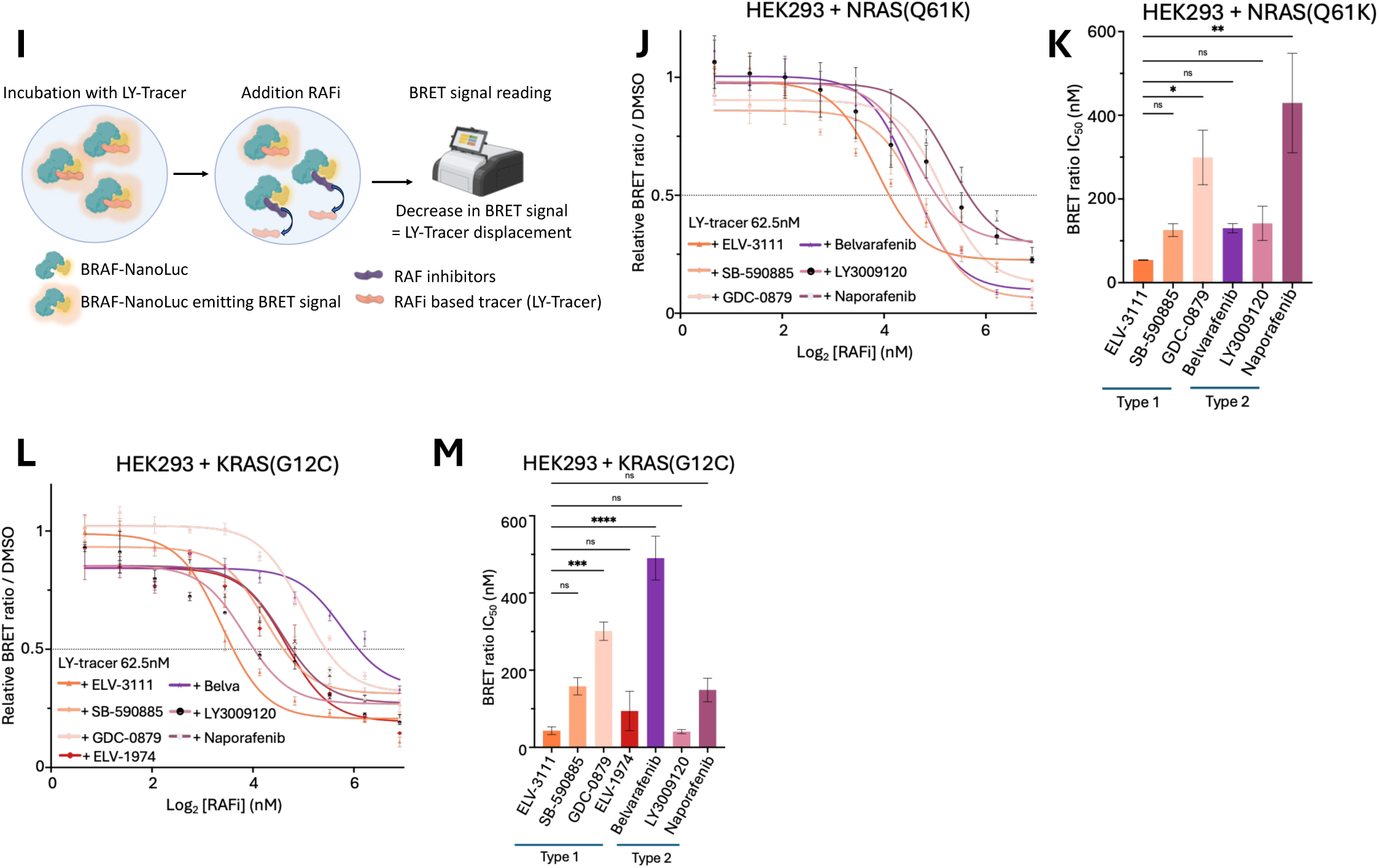
ELV-3111 is a Type 1 RAF inhibitor with broad activity and potent inhibition of both monomeric and dimeric RAF. (**A**) ELV-3111 chemical structure. (**B**) X-ray co-crystal structure of BRAF kinase domain bound to ELV-3111 (orange) or Vemurafenib (blue) at a 2.85Å resolution. (**C**) ELV-3111 was screened at 1µM for activity against 371 kinases with 100µM ATP using DMSO as control. (**D**) NanoLuc tagged A/B/CRAF was expressed in HEK293 cells and treated with K10 (ARAF) or K4 (B/CRAF) tracers and a concentration range of the indicated compounds for 2 hours. BRET ratio was measured and represented as curves (mean values ± SEM*, n* = 3 independent experiments). (**E**) Profiling data for ELV-3111 and other RAFis calculated from dose responses ranging from 1nM to 10,000nM. An in-cell western procedure was used to measure pERK inhibition after 2 hours of treatment. Proliferation assays were performed in the indicated cell lines after 3 (A375 and OCI-AML-2) or 5 (SK-MEL-30) days of treatment. Serum shift was calculated from A375 pERK, except for Naporafenib which was calculated from SK-MEL-30. pERK IC_50_s or GI_50_s were represented as a table (geometric mean, *n* = 3 independent experiments). (**F**) BRAF(V600E) monomeric SK-MEL-239, BRAF(V600E) dimeric resistant SK-MEL-239 C3, NRAS(Q61K) SK-MEL-2 and RAF/RAS(WT) Human Fibroblasts, RPE.1, HPDE and MEFs were treated with increasing concentrations of the indicated RAFis for 6 days. Cell viability was measured and represented as relative viability compared to DMSO-treated controls (mean values ± SEM, *n* = 3 independent experiments). (**G**) BRAF(V600E) monomeric SK-MEL-239, BRAF(V600E) dimeric resistant SK-MEL-239 C3, NRAS(Q61K) SK-MEL-2 and RAF/RAS(WT) Human Fibroblasts, RPE.1, HPDE and MEFs were treated with increasing concentrations of different types of RAFis for 1 hour. Cell lysates were analyzed by immunoblotting with the indicated antibodies to assess MAPK signaling. MEK was used as loading control. (**H**) HeLa cells engineered to express doxycycline-inducible KRAS(Q61K) were treated with doxycycline (500ng/mL) for 48 hours, followed by treatment with increasing concentrations of different types of RAFis for 1 hour. Cell lysates were analyzed by immunoblotting with the indicated antibodies to assess MAPK signaling. MEK was used as loading control. Exposure times were adjusted separately for −Dox and +Dox conditions to visualize treatment effects; baseline signaling is shown in fig. S2, E and F. (**I**) Schematic representation of the *in-cell* BRET target engagement assay. HEK293 cells were co-transfected with BRAF-NanoLuc and NRAS(Q61K) or KRAS(G12C) constructs. Cells were then co-treated with 62.5nM of LY-Tracer and increasing concentrations of different types of RAFis for 2 hours. After treatment, cells were incubated with 20μM Fluorofurimazine, and BRET signal was measured. (**J and L**) Data are presented as relative BRET ratios normalized to DMSO-treated controls (mean values ± SEM, *n* = 3 independent experiments). (**K and M**) BRET ratio IC_50_s values were calculated and represented as bar graphs (mean values ± SEM, *n* = 3 independent experiments, ns = not significant (P>0.05), *P<0.05, **P<0.01, ***P<0.001, ****P<0.0001, ordinary one-way ANOVA).

To determine the binding profile and functional activity of ELV-3111 across RAF isoforms and RAF-driven cellular contexts, we employed a NanoBRET cell-based tracer displacement assay(*18*). ELV-3111 was found to potently bind ARAF, BRAF and CRAF, with similar affinity to the Type 2 RAFi Naporafenib(*19*) and with greater binding of CRAF compared to either the Type 1 RAFi GDC-0879(*20*) or the Type 1.5 RAFi Encorafenib(*13*) (**Fig. 1D and fig. S2A**). ELV-3111, and these same RAFis were then evaluated for their ability to suppress MAPK signaling in both the BRAF(V600E) A375 and NRAS(Q61K) SK-MEL-30 cell lines, in normal cell culture medium as well as in the presence of 100% human serum to model the attenuating effects of plasma protein binding when dosed in patients. ELV-3111 potently suppressed MAPK signaling in both RAF monomer- and dimer-driven contexts, with superior activity compared to all the other RAFis tested (**Fig. 1E**). The antiproliferative activities of these RAFis were assessed in the A375, SK-MEL-30 and OCI-AML-2 cell lines, the latter of which harbors a CRAF fusion gene. ELV-3111 was the only RAFi tested that could inhibit growth of all three cell lines with similar potency (**Fig. 1E**), as Naporafenib has been shown to have reduced potency against monomeric BRAF(V600X)(*21*).

To define the breadth of activity of ELV-3111 in tumor cells, we compared its MAPK pathway and cell growth-suppressing effects to those of other RAFis with various binding modes. Two Type 1 RAFis (ELV-3111 and GDC-0879), one Type 1.5 RAFi (Encorafenib), and two Type 2 RAFis (Naporafenib and LY3009120) were evaluated in cellular models with either monomeric BRAF(V600E) (parental SK-MEL-239), dimeric BRAF(V600E) (clone C3 SK-MEL-239, expressing the splice variant p61BRAF(V600E)), which is known to drive resistance to the approved Type 1.5 RAFis due to p61BRAF(V600E) homodimerization(*22*). SK-MEL-2 cells, signaling via dimeric RAF downstream of mutant RAS (RAS(MUT)), and primary Human Fibroblasts, Human Pancreatic Duct Epithelial cell line (HPDE), RPE.1 cells, and Mouse Embryonic Fibroblasts (MEFs) were used as *in vitro* models of RAS(MUT) and RAS(WT)/BRAF(WT) (representing normal RAS signaling), respectively. Consistent with its binding mode, treatment with ELV-3111 potently inhibited cell growth and MAPK signaling in both monomeric and dimeric BRAF(V600E) cells (**Fig. 1, F and G and fig. S2B**). As expected, the Type 1.5 RAFi Encorafenib exhibited more potent cell growth inhibition in the monomeric model than in the dimeric model (∼720-fold difference). The Type 2 RAFi Naporafenib exhibited greater inhibition in the dimeric model (∼150-fold difference), while LY3009120 displayed equipotent activity in both models, consistent with previous reports(*5, 21, 23*) (**Fig. 1F and fig. S2B**). Further, ELV-3111 suppressed MAPK signaling and cell growth more potently in RAS(MUT) compared to RAS(WT) cells, with RAS(MUT) selectivity ratio of ∼5-fold for growth inhibition and ∼23-fold for MAPK pathway inhibition (**Fig. 1, F and G and fig. S2, C and D**). The potency and selectivity of ELV-3111 exceeded those of the Type 1 RAFi GDC-0879, as well as of the Type 2 RAFis Naporafenib and LY3009120. As expected, the Type 1.5 RAFi Encorafenib was inactive in both RAS(MUT) and RAS(WT) contexts, consistent with its inability to inhibit RAF dimers. These observations correlated well with the MAPK inhibition potency of each RAFi in the cell line models used (**Fig. 1G and fig. S2D**).

Further, we evaluated ELV-3111 together with representative Type 1, Type 2, or Type 1.5 RAFis in an isogenic model with doxycycline-inducible RAS(MUT) expression. As expected, induction of RAS(MUT) to near-endogenous levels enhanced MAPK signaling in both models (**fig. S2, E and F**). All Type 1 and Type 2 RAFis demonstrated preferential MAPK pathway suppression in RAS(MUT) over RAS(WT) cells. ELV-3111 showed a modestly higher degree of separation between these contexts compared with GDC-0879, Naporafenib, or LY3009120. (**Fig. 1H and fig. S2G)**. At concentrations within a therapeutic range, RAS(MUT) induction did not significantly enhance MAPK inhibition with Encorafenib. A slight differential effect was only observed at supraphysiological concentrations (∼10μM), consistent with the inability of Type 1.5 inhibitors to potently inhibit RAF dimers.

To further substantiate these findings, we compared the activity of ELV-3111 to representative Type 1.5 (Encorafenib) and Type 2 (Naporafenib) RAFis, as well as a MEKi (Binimetinib) in Ba/F3 cells engineered to express distinct classes of BRAF mutations. In this system, ELV-3111 exhibited superior and broader inhibitory activity on cell viability across both monomeric and dimeric RAF-driven signaling contexts (**fig. S3, A and B)**, indicating enhanced functional coverage compared to existing RAFi classes. We further used COLO 201 cells endogenously expressing BRAF(V600E) and engineered them to also express oncogenic RAS(MUT) to model acquired resistance to Type 1.5 RAFis mediated by RAS-driven RAF dimerization(*24, 25*). While the Type 1.5 RAFi Encorafenib exhibited reduced potency in these RAF dimer-dependent models, ELV-3111 remained highly effective in suppressing growth and MAPK signaling of BRAF(MUT)/RAS(MUT) cells **(fig. S3, C to F**).

We finally evaluated ELV-3111 across a broader panel of tumor cell lines, including RAS(WT) and various RAS(MUT) and BRAF(MUT) models. ELV-3111 demonstrated high potency in cells driven by Class 1, Class 2, and Class 3 BRAF mutants, with GI_50_ and IC_50_ values under 100nM, and was also highly effective against multiple NRAS- and KRAS-mutant cell lines and cell lines harboring dual mutations in both BRAF and NRAS (**fig. S4, A to F and Table S3**).

Both Type 1 and Type 2 kinase inhibitors bind in the αC-helix-IN mode(*4*), which is associated with the active dimeric RAF conformation. We thus asked whether the high potency of ELV-3111 in RAS(MUT) tumor cells is the result of strong inhibitor binding to the highly active RAF conformation in RAS(MUT) cells. To this end, we developed a live-cell NanoBRET BRAF engagement assay(*18, 26*) to monitor target engagement of RAF inhibitors (**Fig. 1I**). HEK293 cells were co-transfected with BRAF-NanoLuc with or without NRAS(Q61K) and KRAS(G12C) for 24 hours, followed by treatment with increasing concentrations of a fluorescent tracer that we designed and synthesized based on the potent RAFi LY3009120 (LY-Tracer – **fig. S5A**). The apparent dissociation constant (LY-Tracer K_D,app_) was first determined to select the LY-Tracer concentration for subsequent competition assays (∼50% of RAF-binding saturation) (**fig. S5B**). Ectopic RAS(MUT) expression resulted in a significant increase in NanoBRET signal by LY-Tracer, at much higher levels compared to non-transfected cells, consistent with active RAS(MUT) promoting the active conformation of RAF that enhances RAFi binding. ELV-3111 was consistently highly potent in RAS(MUT)-expressing cells, demonstrating a ∼2-fold greater potency than LY3009120, 4-fold greater than Naporafenib, and 5-fold greater than Belvarafenib - all Type 2 RAFis (**Fig. 1, J to M and fig. S5, C to E**). Thus, ELV-3111 exhibits high activity in RAS(MUT) cells, through enhanced target engagement and potent inhibition of fully activated RAF.

Taken together, these data demonstrate that ELV-3111 is broadly active across many different classes of MAPK alterations, including settings of known resistance to approved Type 1.5 RAFis, and maintains activity in both the monomer and dimer RAF settings, with a measurable therapeutic window in dimeric RAF-driven, including RAS(MUT) tumors, over normal cells.

### ELV-3111 potently suppresses MAPK in BRAF(MUT) and RAS(MUT) tumor models while inducing MAPK hyperactivation in normal tissues as drug concentrations decline *in vivo*

To assess MAPK pathway inhibition by ELV-3111 *in vivo*, we carried a pharmacodynamic/pharmacokinetic study (PK/PD) in mice engrafted with RAS(MUT) (SK-MEL-30 melanoma) xenograft tumors (**Fig. 2A**). As expected, after a single dose of ELV-3111, initial MAPK inhibition was followed by recovery in both tumor and normal tissues, consistent with relief of negative feedback and adaptive response to MAPK-targeting inhibitors previously described by us and others(*27–30*). All treatments potently inhibited the pathway with a modest MAPK recovery in RAS(MUT) tumors (**Fig. 2B**). Type 2 RAFi Naporafenib and the FDA-approved MEK inhibitor (MEKi) Binimetinib, as single treatments, resulted in MAPK pathway inhibition in normal tissues and recovery close to baseline, as expected (**Fig. 2C and fig. S6A**). Surprisingly, treatment with ELV-3111 resulted in marked MAPK hyperactivation above basal levels in normal tissue (∼ 3 and ∼2 fold above baseline in lung and kidney, respectively - **Fig. 2C and fig. S6A)**. This hyperactivation emerges as ELV-3111 plasma concentration declined following peak exposure (**Fig. 2D**), suggesting a relationship between drug concentration and MAPK signaling dynamics. Naporafenib, despite exhibiting a similar PK profile and *in vivo* half-life (**Fig. 2D**), did not induce MAPK hyperactivation. Thus, ELV-3111 effectively suppressed MAPK signaling in RAS(MUT) tumors, similar to Type 2 RAFis, but paradoxically hyperactivated MAPK signaling in normal tissues. This phenomenon is reminiscent of that observed with first-generation Type 1.5 RAFis, and indicates an obvious opportunity for substantially expanding the therapeutic index using this RAFi. We next assessed the combination of either ELV-3111 or Naporafenib with Binimetinib in the PK/PD study. The combination of Type 2 RAFi with Binimetinib enhanced MAPK inhibition in both tumor and normal cells, consistent with the narrow therapeutic window of these drugs. ELV-3111 combined with Binimetinib strongly inhibited MAPK in the tumor; however, in normal cells, the opposing effects of the two drugs restored MAPK levels close to normal (**Fig. 2, B and C and fig. S6A**). These data indicate that paradoxical MAPK hyperactivation by Type 1 RAFi selectively in normal tissue enables expanding the therapeutic index through combinations with MAPK inhibitors.

**Fig. 2.**
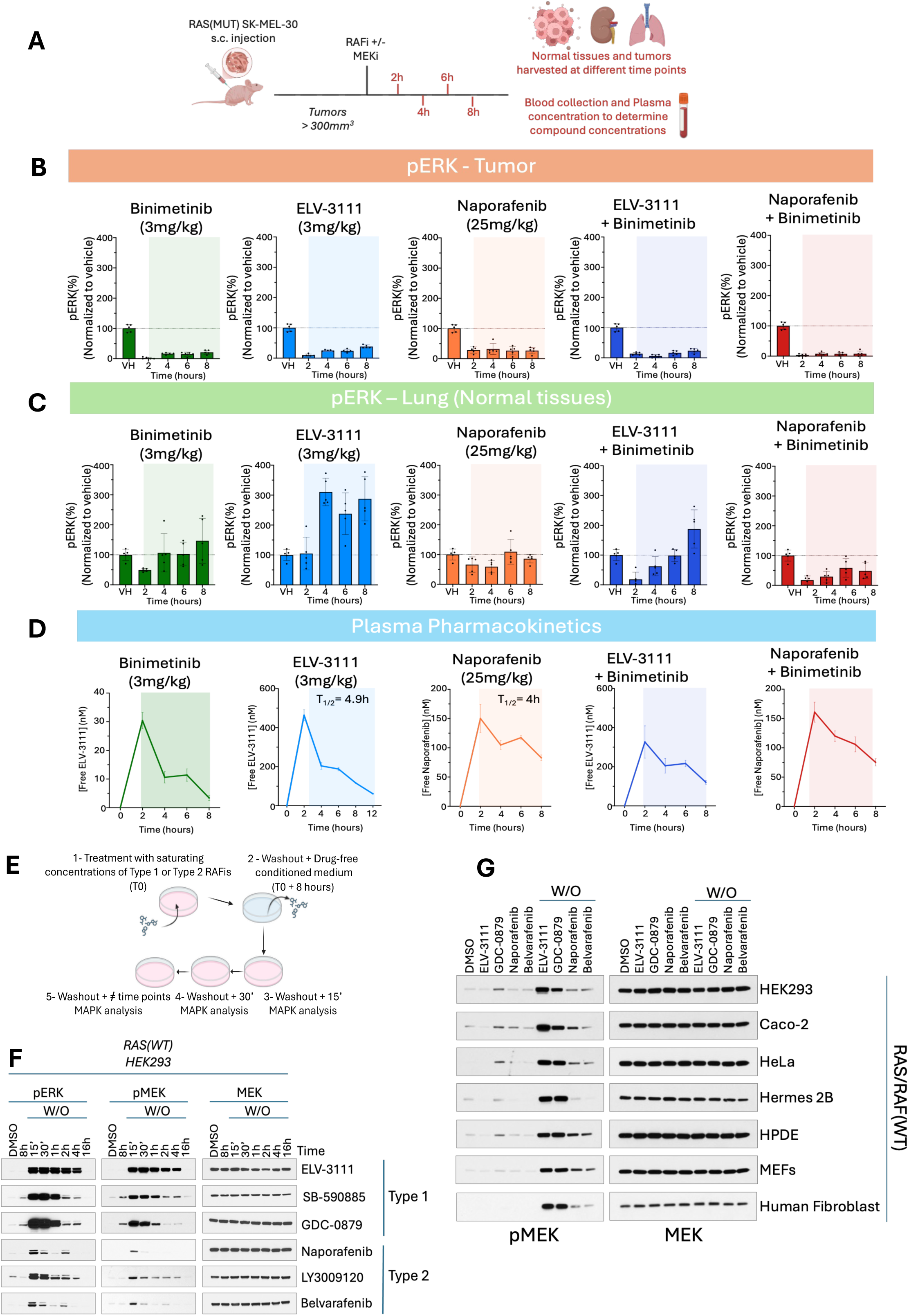
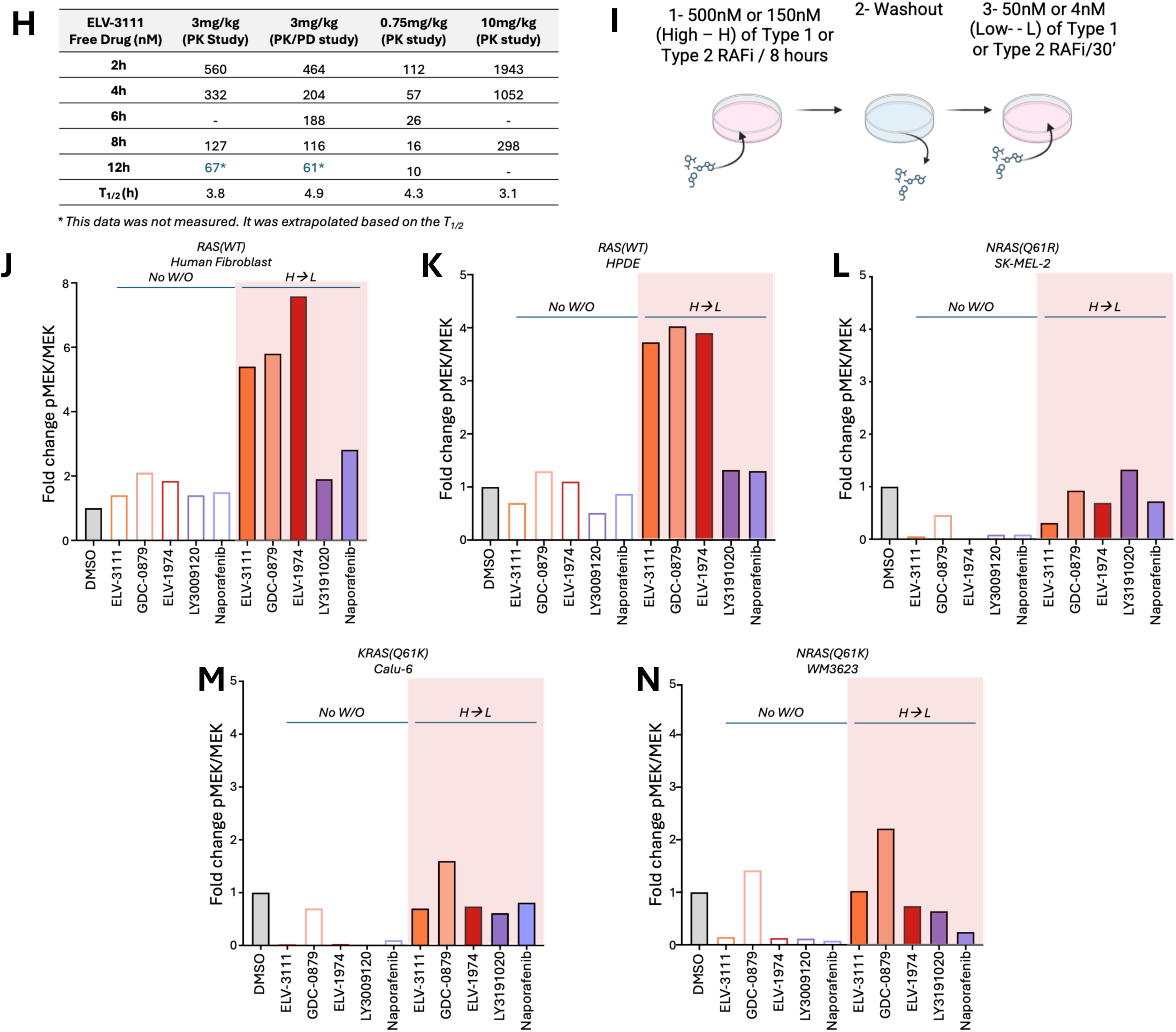
ELV-3111 potently suppresses MAPK in BRAF(MUT) and RAS(MUT) tumor models while inducing MAPK hyperactivation in normal tissue, *in vivo*. (**A**) Schematic representation of the experimental design used to assess *in vivo* pharmacodynamic/pharmacokinetic effects of ELV-3111 (Type 1 RAFi) or Naporafenib (Type 2 RAFi) in combination or not with Binimetinib (MEKi). SK-MEL-30 tumor xenograft-bearing mice were dosed via oral gavage with a single dose of 3mg/kg Binimetinib, 3mg/kg ELV-3111, 25mg/kg Naporafenib, or combinations of Binimetinib + ELV-3111 or Binimetinib + Naporafenib. (**B**) Tumor and (**C**) lung tissues were collected after 2, 4, 6 or 8 hours, lysed, and analyzed by immunoblotting for pERK to assess MAPK pathway activity. Data are represented as the mean relative pERK levels normalized to vehicle-treated controls (*n* = 5 mice per group). (**D**) Plasma concentration of free RAFis was measured 2, 4, 6 or 8 hours after dosing. (**E**) Schematic representation of the experimental design used for the complete drug washout time-course studies. (**F**) RAS(WT) HEK293 cells were treated with 1μM of either Type 1 RAFis (ELV-3111, SB-590885), Type 2 RAFis (Naporafenib, LY3009120, Belvarafenib) or 10μM of the Type 1 RAFi GDC-0879. Following treatment, cells were washed and incubated in drug-free conditioned medium for the indicated time points. Cell lysates were analyzed by immunoblotting to assess MAPK signaling. MEK was used as loading control. (**G**) A panel of RAS(WT) cell lines (HEK293, Caco-2, HeLa, Hermes 2B, HPDE, MEFs and Human Fibroblast) was treated with 1μM of either Type 1 RAFi ELV-3111, Type 2 RAFis (Naporafenib or Belvarafenib), or 10μM of the Type 1 RAFi GDC-0879 for 8 hours. Following treatment, cells were washed and incubated in drug-free conditioned medium for 15 or 30 minutes, depending on the cell lines. Cell lysates were analyzed by immunoblotting to assess MAPK signaling. MEK was used as loading control. (**H**) Concentration of ELV-3111 was measured and adjusted for known plasma protein binding to determine the free drug concentration (nM) at the indicated time points from 3 pharmacokinetic studies in mice (*n* = 3 animals/group), or the PK/PD study in (A). (**I**) Schematic representation of the experimental design for the gradual drug washout studies. (**J**) RAS(WT) Human Fibroblast, (**K**) RAS(WT) HPDE, (**L**) NRAS(Q61R) SK-MEL-2, (**M**) KRAS(Q61K) Calu-6, or (**N**) NRAS(Q61K) WM3623 cells were treated with 500nM (High – H) of Type 1 RAFis (ELV-3111, GDC-0879, ELV-1974) or Type 2 RAFis (LY3009120, Naporafenib) for 8 hours. Following treatment, cells were washed and incubated with 50nM (Low – L) of Type 1 or Type 2 RAFis for 15 or 30 minutes depending on the cell line. Cell lysates were analyzed by immunoblotting (pMEK, MEK) to assess MAPK signaling. Ratios of phosphorylated MEK to MEK for each sample were calculated and represented as fold change pMEK bar graphs compared to DMSO-controls.

Unlike the paradoxical MAPK hyperactivation induced by Type 1.5 RAFis that is evident both *in vitro* and *in vivo*(*8–10*), MAPK hyperactivation by ELV-3111 appears more robust *in vivo*. *In vitro* studies with a single fixed concentration of compound over time do not faithfully replicate the dynamic nature of an *in vivo* pharmacokinetic (PK) profile, where compound concentrations spike and then decrease with inhibitor clearance (**Fig. 2D**). To better model the *in vivo* setting, we first treated cells with different RAFis followed by washout to simulate drug clearance. First, RAS(WT) HEK293 cells, as a model of normal cells, were treated with either Type 1 RAFis (ELV-3111, GDC-0879 or SB-590885), or Type 2 RAFis (including Naporafenib, used in the PK/PD studies) at saturating concentrations for 8 hours, followed by washout (**Fig. 2E**). Treatment with ELV-3111 or any Type 1 RAFi, followed by washout, resulted in robust MAPK hyperactivation (**Fig. 2F**), indicating that this is a shared property of Type 1 RAFis. In contrast, treatment with Type 2 RAFis resulted in a significantly lower MAPK rebound, consistent with the Naporafenib profile *in vivo*. To further extend these observations, we treated an expanded panel of RAS(WT) cells with two Type 1 RAFis (ELV-3111, GDC-0879) or two Type 2 RAFis (Naporafenib, Belvarafenib), followed by washout. Consistently, upon wash-out, each Type 1 RAFi induced pronounced MAPK hyperactivation, whereas MAPK rebound after Type 2 RAFi treatment was minimal (**Fig**. **2G and fig****. S6B**).

To simulate more faithfully the gradual reduction of drug concentration observed *in vivo*, we established two settings of high and low ELV-3111 concentrations - Cmax of 150 nM or 500 nM and Cmin of 4 nM or 50 nM - based on free drug concentrations we observed in pharmacokinetic studies in mice (**Fig. 2H**). RAS(WT) cells and RAS(MUT) were initially treated at a peak concentration (150 nM or 500 nM) with a panel of Type 1 RAFis, including ELV-3111, GDC-0879 and ELV-1974, a Type 1 RAFi sharing a closely related core scaffold with ELV-3111 (**fig. S6C**), or with Type 2 RAFis LY3009120 or Naporafenib. Following washout, cells were re-exposed to the corresponding low concentration (4 nM or 50 nM, respectively – **Fig. 2I**). Under these conditions, Type 1 RAFis induced significantly higher MAPK hyperactivation than Type 2 RAFis, selectively in RAS(WT) cells but not in RAS(MUT) cells (**Fig. 2, J to N and fig. S6, D and E**). GDC-0879 showed modest hyperactivation in RAS(MUT) cells(*10*), consistent with its low potency and incomplete suppression of primed RAF at the concentrations used, as shown for αC-IN RAF inhibitors(*5*). To validate these findings by an orthogonal approach, we tested the Type 1 RAFi ELV-3111 and the Type 2 RAFi Naporafenib in two RAS(WT) models and one RAS(MUT) model, using a TR-FRET assay to measure pERK. Consistent with the preceding results, ELV-3111 induced significantly greater MAPK hyperactivation than Naporafenib, selectively in RAS(WT) cells (**fig. S6F**).

Together, these results support a model in which MAPK hyperactivation in normal cells following ELV-3111 treatment is a property of Type 1 RAFis that separates them from Type 2 RAFis and is the consequence of declining drug concentrations *in vivo*.

### Type 1 RAFi-induced MAPK hyperactivation is RAS-dependent

To understand the mechanism underlying the Type 1 RAFi-mediated MAPK hyperactivation phenotype in normal cells, we first investigated the potential role of RAS(WT) activity. We have shown previously that RAS activity is essential for MAPK paradoxical activation by Type 1.5 RAFis(*5*). To test whether this is also the case for Type 1 RAFis, we first used a SHP2 inhibitor (TNO155)(*31*) and the EGFR inhibitor Gefitinib, both of which suppress RAS(WT) activity. In RAS(WT) cells, high concentrations of either inhibitor in combination with ELV-3111 markedly suppressed MAPK signaling and prevented hyperactivation induced by ELV-3111 after washout (**Fig. 3, A and B**). We further validated this dependency using HEK293 cells in which all three RAS paralogs have been knocked out via CRISPR (“RAS-less cells”)(*32*). Consistent with our hypothesis, ELV-3111-induced MAPK hyperactivation was abolished in RAS-less cells after washout and was fully restored upon ectopic expression of RAS(WT) (**Fig. 3C**).

**Fig. 3.**
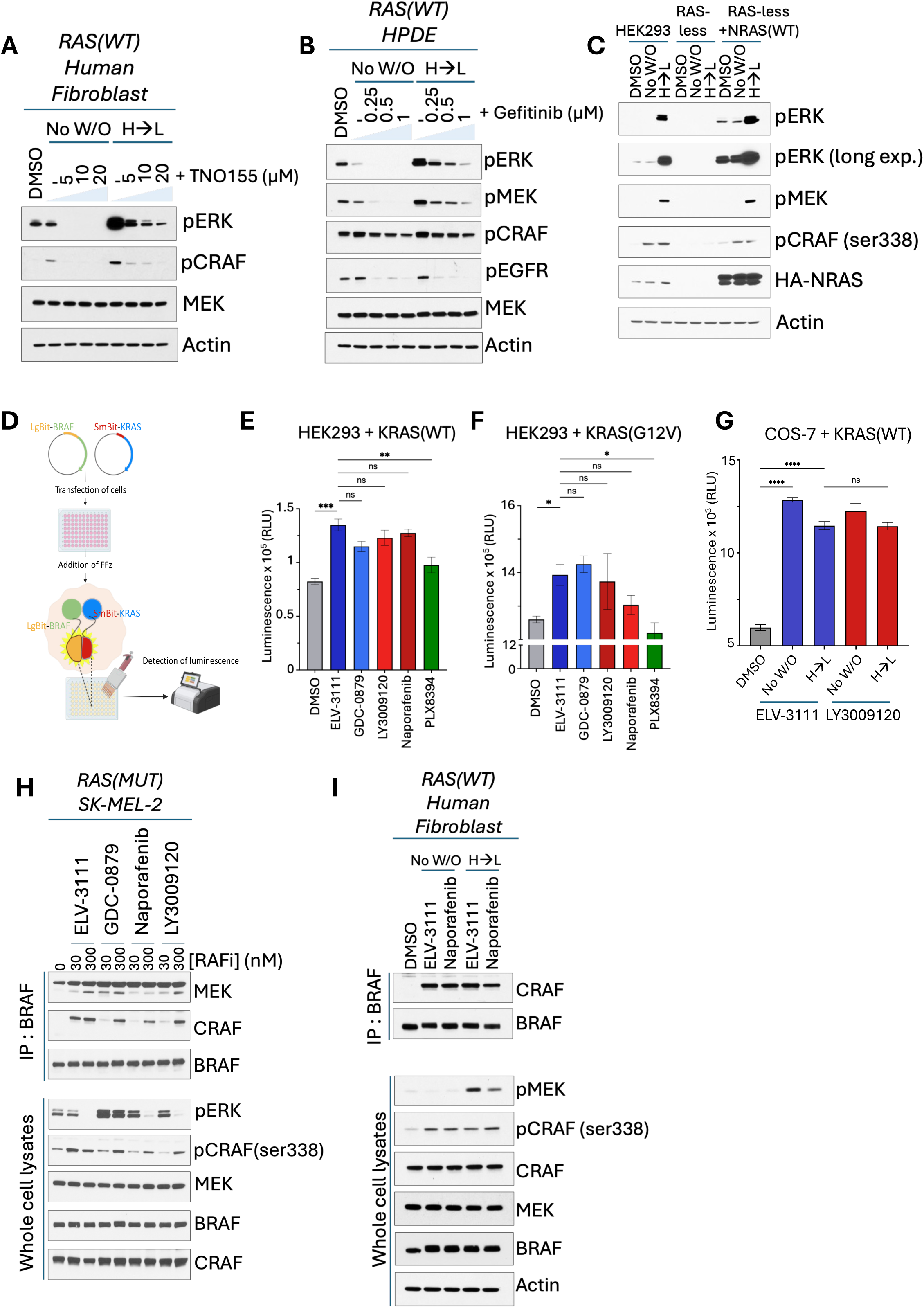
Type 1 RAFi-induced MAPK hyperactivation is RAS-dependent. (**A**) Human Fibroblast cells were treated with 500nM (High – H) of ELV-3111 in combination or not with increasing concentrations of SHP2 inhibitor (TNO155). After treatment, cells were washed and incubated with 50nM of ELV-3111 (Low – L) and the same concentrations of SHP2 inhibitor. Cell lysates were immunoblotted to assess MAPK signaling. MEK and actin were used as loading controls. (**B**) HPDE cells were treated with 500nM (High – H) of Type 1 ELV-3111 in combination or not with increasing concentrations of EGFR inhibitor (Gefitinib). After treatment, cells were washed and incubated with 50nM of ELV-3111 (Low – L) and the same concentrations of EGFR inhibitor. Cell lysates were subjected to immunoblotting to assess MAPK signaling. MEK and actin were used as loading controls. (**C**) HEK293, HEK293 RAS-less or HEK293 RAS-less cells ectopically expressing NRAS(WT) were treated with 500 nM (High – H) of Type 1 ELV-3111. Following treatment, cells were washed and incubated with 50 nM (Low – L) of ELV-3111. Cell lysates were immunoblotted with the indicated antibodies to assess MAPK signaling and RAS expression. Actin was used as loading control. (**D**) Schematic representation of the NanoBit assay used to assess BRAF-KRAS interaction in live-cells. HEK293 cells were co-transfected with LgBit-BRAF and either (**E**) SmBit-KRAS(WT) or (**F**) SmBit-KRAS(G12V) constructs for 24 hours. Cells were then treated with 300nM of different Type 1 RAFis (ELV-3111, GDC-0879), Type 2 RAFis (Naporafenib, LY3009120) or Type 1.5 Paradox Breaker (PLX8394). Immediately after treatment, cells were incubated with 20μM of Fluorofurimazine, and luminescence signal was measured over time. Data are presented as luminescence after 60 minutes of treatment/plate reading and represented as bar graphs (luminescence mean values ± SEM, *n* = 3 independent experiments; ns = not significant (P>0.05), *P<0.05, **P<0.01, ***P<0.001, ordinary one-way ANOVA). (**G**) COS-7 cells were co-transfected with LgBit-BRAF and SmBit-KRAS(WT) constructs for 24 hours. Cells were then treated with 500nM (High – H) of Type 1 RAFi ELV-3111 or Type 2 RAFi LY3009120 for 2 hours. Following treatment, cells were washed, treated with 50nM (Low – L) of ELV-3111 or LY3009120, and directly incubated with 20μM of Fluorofurimazine. Luminescent signal was measured over time. Data are presented as luminescence after 60 minutes of treatment/plate reading and represented as bar graphs (mean luminescence values ± SEM, *n* = 3 independent experiments; ns = not significant (P>0.05), ****P<0.0001, ordinary one-way ANOVA). (H) NRAS(Q61R) SK-MEL-2 cells were treated with different concentrations (0.03, 0.3μM) of the indicated RAFis for 1 hour. BRAF was immunoprecipitated and immunoblotted with the indicated antibodies to assess BRAF-CRAF dimerization and BRAF-MEK interaction. Whole cell lysates were immunoblotted with the indicated antibodies to assess MAPK signaling. (**I**) RAS(WT) Human Fibroblast cells were treated with 500nM (High – H) of Type 1 RAFi ELV-3111 or Type 2 RAFi Naporafenib. Following treatment, cells were washed and treated with 50nM (Low – L) of either ELV-3111 or Naporafenib. BRAF proteins were immunoprecipitated and immunoblotted with the indicated antibodies to assess BRAF-CRAF dimerization. Whole cell lysates were immunoblotted with the indicated antibodies to assess MAPK signaling.

Both Type 1 and Type 2 RAFis are potent RAF dimer inhibitors. However, Type 1 RAFis promoted significantly higher levels of paradoxical MAPK hyperactivation in normal cells compared to Type 2 RAFis. To understand the basis of the difference, we compared their biochemical properties in more detail. First, we assessed formation of the RAS-RAF complex by employing a NanoBiT complementation proximity assay(*33*). In this system, RAS and BRAF are tagged with the NanoLuc subunits SmBiT and LgBiT, respectively, and expressed in HEK293 cells (**fig. S7A)**. Luminescence is generated upon complex assembly, providing a quantitative readout of protein-protein interactions in live-cells(*33*) (**Fig. 3D**). We measured RAS-BRAF interaction after treatment with Type 1 or Type 2 RAFis, as well as a Type 1.5 RAFi that does not induce paradoxical RAF/MAPK activation (Plixorafenib – PLX8394)(*34*) as a negative control. As expected, and consistent with previous reports(*5, 35*), both Type 1 and Type 2 RAFis enhanced RAS-BRAF interaction to a similar extent, whereas Plixorafenib did not (**Fig. 3, E and F**). To mimic drug clearance *in vivo*, we repeated the experiment by treating with RAFis, followed by drug washout. Treatment with the Type 1 RAFi ELV-3111 or the Type 2 RAFi LY3009120, followed by washout, resulted in a similar induction of the RAS-BRAF interaction in RAS(WT) setting (**Fig. 3G**). Thus, the induction of the RAS-BRAF interaction is a property shared by Type 1 and Type 2 RAFis and cannot account for their different phenotype *in vivo*.

To further characterize RAFi-induced signaling complexes, we examined the BRAF-CRAF and BRAF-MEK complex formation following treatment with different RAFis. Both Type 1 and Type 2 RAFis induced RAF transition to its active state, also known as RAFi-induced “priming”(*5, 10*), evidenced by an increase in S338 phosphorylation of CRAF, BRAF-CRAF dimerization, and BRAF-MEK complex formation at comparable levels in both RAS(MUT) (**Fig. 3H and fig. S7, B and C**) and RAS(WT) Human Fibroblast cells (**Fig. 3I**), after washout. ELV-3111 also promoted the BRAF-CRAF complexes at much lower concentrations (**Fig. 3H)**, consistent with its higher RAF binding potency. We also profiled the formation of BRAF or CRAF homodimers or BRAF-CRAF heterodimers using NanoBRET technology and found ELV-3111 to be distinct from other known Type 1 RAFis in its ability to drive CRAF homodimers to a similar extent as Naporafenib (**fig. S7D**). Thus, induction of the RAF signaling complexes is similar between Type 1 and Type 2 RAFis, it is therefore unlikely to cause their different phenotype *in vivo*. Finally, to exclude the possibility that Type 1 RAFi induces MAPK hyperactivation through higher RAS-GTP levels resulting from RTK-RAS feedback reactivation, we treated RAS(WT) HPDE cells with low concentration of ELV-3111 (selected to avoid MAPK pathway inhibition and thereby limit relief of ERK-dependent negative feedback) or with an ERK inhibitor as a positive control, either alone or in combination **(fig. S7, E and F)**. As expected, ERK inhibition caused classical feedback relief, resulting in increased RAS activity. In contrast, ELV-3111 washout induced MAPK hyperactivation without increasing RAS activity. Notably, combining ELV-3111 with the ERK inhibitor did not further increase RAS activity compared with ERK inhibition alone. Together, these findings demonstrate that ELV-3111-mediated MAPK hyperactivation is mechanistically distinct from classical RTK–RAS feedback reactivation.

### Type 1 RAFis exhibit over an order-of-magnitude faster dissociation from RAF relative to Type 2 RAFis

As both Type 1 and Type 2 RAFis promoted the active RAF conformation and similar RAF signaling complexes, we next hypothesized that differences in inhibitor off-rate kinetics may promote inhibitor dissociation and activated RAF activity following Type 1, but not Type 2, RAFi withdrawal. To test this, we characterized the binding kinetics of two Type 1 RAFis (ELV-3111, GDC-0879) and two Type 2 RAFis (Naporafenib, LY3009120), using an *in vitro* TR-FRET binding assay based on competition with a fluorescent probe targeting purified activated BRAF (**Fig. 4A**). Probe occupancy occurred much more rapidly for Type 1 RAFis compared to Type 2 RAFis, indicating a faster dissociation rate (**Fig. 4B**). Calculation of dissociation rate constant (k_2_) and residence times (τ) revealed that ELV-3111 and GDC-0879 exhibited markedly faster - by about an order of magnitude - off-rates and thus possess substantially shorter residence times than Type 2 RAFis (**fig. S8, A and B**).

**Fig. 4.**
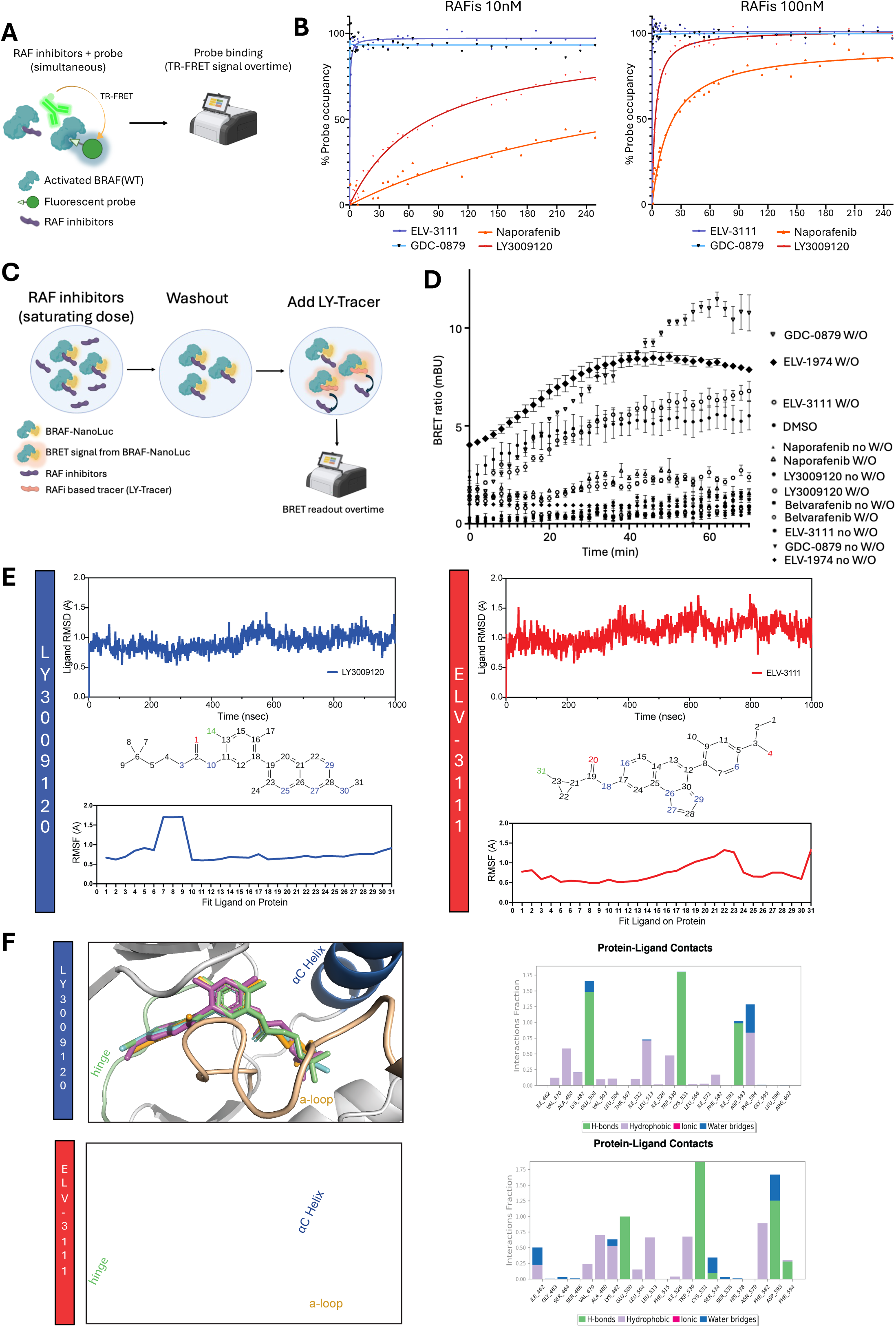
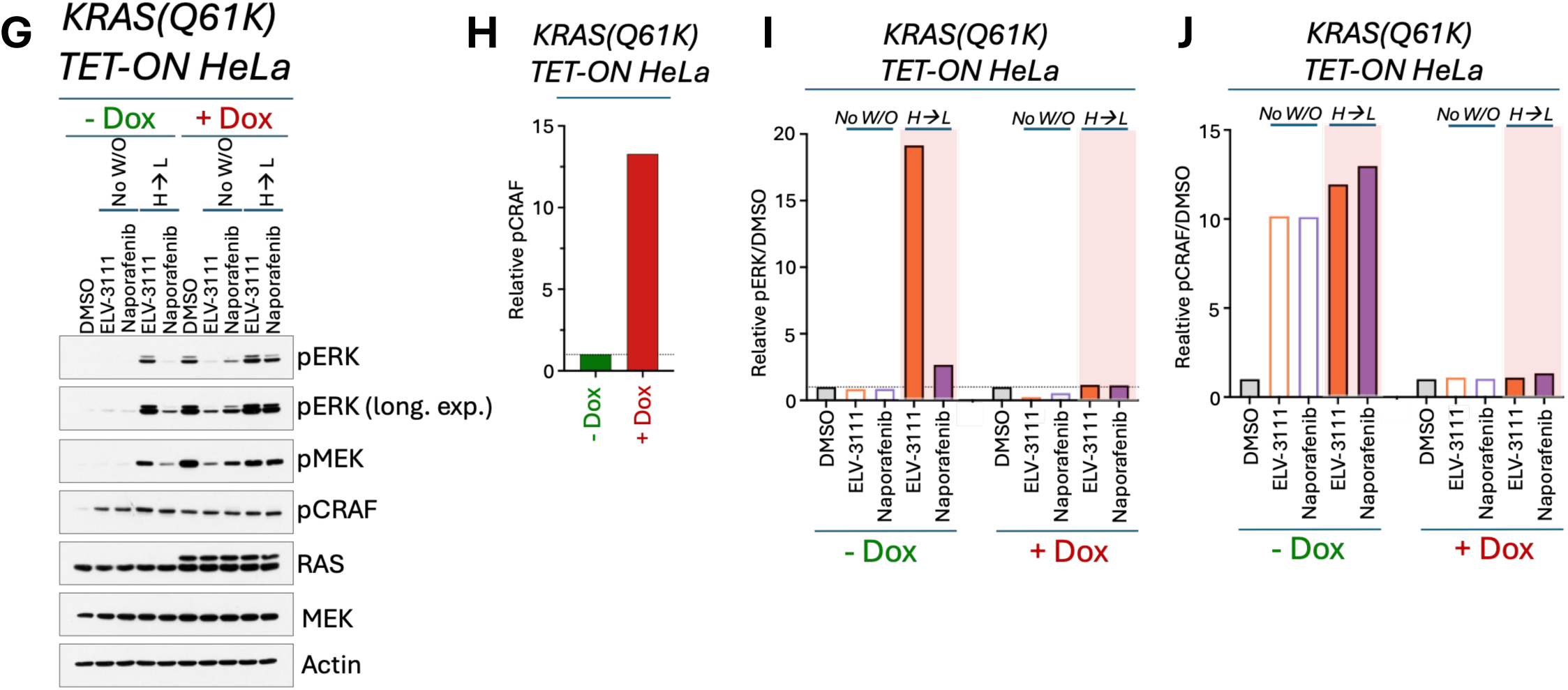
Type 1 RAFis exhibit over an order-of-magnitude faster dissociation from RAF relative to Type 2 RAFis. (**A**) Schematic representation of the *in vitro* TR-FRET binding assay based on competition with a fluorescent probe targeting purified activated BRAF. (**B**) The binding kinetics of Type 1 RAFis (ELV-3111, GDC-0879) or Type 2 RAFis (Naporafenib, LY3009120) were characterized at two concentrations (10nM and 100nM) against 0.3nM of activated BRAF(WT) *in vitro*. % of probe occupancy over time was represented as curves (**C**) Schematic representation of the NanoBRET assay used to assess RAFi residence time in live cells. (**D**) COS-7 cells were transfected with BRAF-NanoLuc for 24 hours. Cells were then treated for 2 hours with saturating concentrations of the indicated Type 1 (ELV-3111, GDC-0879, ELV-1974) or Type 2 (Naporafenib, LY3009120, Belvarafenib) RAFis. After treatment, cells were washed, treated with 62.5nM of LY-Tracer and incubated with 20μM of Fluorofurimazine. BRET ratio signals were measured over time using a plate reader (mean values ± SEM, *n* = 3 independent experiments). (**E**) Ligand RMSD plots showing the fluctuation of LY3009120 (Top -blue) and ELV-3111 (Top - red) over 1 μs MD simulations. (Bottom) Chemical structures of LY3009120 and ELV-3111 with atomic numbering alongside their respective ligand RMSF profiles, highlighting atomic fluctuations within the binding pocket. (**F**) Overlays of ligand conformations extracted from four time points (200 ns, 400 ns, 600 ns, and 800 ns) from the 1 μs MD trajectory (**Left**). Two-dimensional ligand interaction diagrams summarizing the key residues mediating contacts with LY3009120 and ELV-3111 (Right). <u>Note</u>: Residue numbering is shifted by one position in the BRAF sequence relative to the crystal structure/MD output. For example, Glu500 in the figure corresponds to Glu501 in the canonical BRAF sequence. (**G**) HeLa cells engineered to express doxycycline-inducible KRAS(Q61K) were treated with doxycycline (500ng/mL) for 48 hours, followed by treatment with 500nM (High – H) of Type 1 RAFi (ELV-3111) or Type 2 RAFi (Naporafenib) for 8 hours. Following treatment, cells were washed and treated with 50nM (Low – L) of either ELV-3111 or Naporafenib. Cell lysates were analyzed by immunoblotting with the indicated antibodies to assess MAPK signaling. MEK and actin were used as loading controls. (**H**) pCRAF levels at baseline and after or without doxycycline induction were quantified from (G) and represented as a bar graph. (**I**) pERK and (**J**) pCRAF levels following RAFi treatment were quantified from the immunoblots shown in (G), normalized to DMSO control, and represented as bar graphs.

To validate these findings in live cells, we developed a NanoBRET-based RAFi competition assay(*18*). NanoLuc-BRAF was ectopically expressed and treated with saturating concentrations of Type 1 (ELV-3111, GDC-0879, and ELV-1974) or Type 2 (LY3009120, Naporafenib, and Belvarafenib) RAFis, followed by washout and addition of LY-Tracer (**Fig. 4C**). We found that the signal after Type 2 RAFis treatment with and without washout remained very low, indicating prolonged binding of RAFis to RAF after washout. In contrast, the signal after ELV-3111, ELV-1974 or GDC-0879 treatments remained low without washout, but dramatically increased after washout, indicating rapid inhibitor dissociation (**Fig. 4D and fig. S8C**). Notably, the signal after washing out either Type 1 RAFi examined was even higher than basal (DMSO-treated), consistent with prior RAF engagement by Type 1 RAFi enhancing LY-Tracer binding, due to induction of the RAF active conformation.

To better understand the structural basis of the difference in the rate of dissociation between Type 1 (ELV-3111 or GDC-0879) and Type 2 (LY3009120 or Sorafenib) RAFis, we carried out molecular dynamics (MD) simulations using their respective crystal structures bound to BRAF. Type 2 RAFis exhibited a lower ligand RMSD throughout the simulation compared to Type 1 RAFis, consistent with a more stable binding conformation (**Fig. 4E and fig. S8, D and E**). Analysis of ligand flexibility showed higher RMSF values for ELV-3111, particularly in solvent-exposed area near the hinge, where the fluorocyclopropane amide moiety adopted dynamic conformations (**Fig. 4, E and F**). While ELV-3111 and LY3009120 inhibitors formed comparable hydrophobic interactions within the ATP-binding cleft and maintained hydrogen bonds with conserved residues Cys531, Asp593 and Glu500, LY3009120’s urea group enabled persistent bifurcated hydrogen bonds with Glu500 over the course of the simulation (**Fig. 4F and fig. S8F**). Thus, the urea-mediated interactions and reduced flexibility of LY3009120 likely confer enhanced stabilization of the inhibitor–kinase complex consistent with its slower dissociation, whereas the increased conformational mobility of ELV-3111 is consistent with its faster dissociation kinetics.

To validate the priming requirement for Type 1 RAFi-mediated MAPK hyperactivation in a single background, we used doxycycline-inducible KRAS(Q61K) systems (**Fig. 4, G to J; fig. S9, A to C**). KRAS(MUT)-induction raises baseline MAPK signaling and pCRAF, indicating enhanced priming. Without doxycycline (RAS(WT)), Type 1 RAFi ELV-3111 and both Type 2 RAFis LY3009120 and Naporafenib increased pCRAF, but only ELV-3111 produced strong MAPK hyperactivation after washout. With KRAS(Q61K) induced, baseline pCRAF was already elevated; neither inhibitor further increased it, and no hyperactivation occurred. Hyperactivation therefore requires induction of priming in a non-primed background and is attenuated where RAF is already primed by oncogenic RAS. Further, we expressed dimerization-deficient RAF mutants (CRAF(R401H) or BRAF(R509H)) or RAF(WT) in HEK293 and assess MAPK hyperactivation after ELV-3111 treatment and washout.

ELV-3111-induced MAPK activation was lower in RAF(MUT) settings compared to wild-type RAF, shown by lower pERK and pMEK. Induction of pCRAF by ELV-3111 was also reduced in the dimerization-deficient mutants, indicating impaired priming (**fig. S9D**). RAF dimerization is therefore required for inhibitor-induced priming and the subsequent hyperactivation, and disrupting it blocks the effect.

Collectively, these data support a model in which both Type 1 and Type 2 RAFis promote RAF association with RAS, RAF dimerization, and transition into the active conformation. However, the rapid dissociation of Type 1 RAFis from activated RAF generates partially or fully drug-free RAF dimers capable of hyperactivating MAPK signaling in normal cells. In RAS(MUT) cells, RAF is already primed, and Type 1 RAFi suppresses RAF and MAPK signaling (**fig. S9E**).

### ELV-3111 - based MAPK targeting combinations are highly potent in BRAF(MUT) and RAS(MUT) tumors and well tolerated *in vivo*

MAPK hyperactivation in normal tissues by Type 1 RAFis indicates that MAPK-targeting vertical combinations would enhance antitumor control and mitigate toxicities by restoring basal levels of MAPK signaling in normal tissue. To evaluate the therapeutic potential of combining MAPK inhibitors with ELV-3111 in BRAF(MUT) or RAS(MUT) tumors, we first assessed the combination of ELV-3111 with the MEKi Binimetinib *in vitro*. The ELV-3111+Binimetinib combination consistently resulted in superior viability inhibition in KRAS(MUT) cell lines compared to treatment with either inhibitor alone (**Fig. 5, A and B**). To investigate the mechanistic basis underlying the enhanced efficacy of the ELV-3111 + Binimetinib combination compared to either agent alone *in vitro*, we assessed MAPK signaling dynamics and apoptotic responses at both early (1 hour) and later (8, 24 or 48 hours) time points. While ELV-3111 or Binimetinib alone induced only transient suppression of pERK, followed by partial rebound of MAPK signaling, the combination induced a more pronounced and sustained inhibition of the MAPK pathway, with minimal or no evidence of pathway reactivation. Consistent with these signaling dynamics, prolonged combination treatment resulted in stronger suppression of Cyclin D1, and greater induction of apoptosis as evidenced by the increased levels of cleaved PARP and cleaved caspase-3/7 activity, compared to single-agent treatments (**Fig. 5, C and D, and fig. S10, A to F**). Furthermore, ELV-3111 demonstrated synergism when combined with Binimetinib in a dose range around its single-agent GI_50_ in both BRAF- and NRAS-mutant cell lines (**fig. S10G**).

**Fig. 5.**
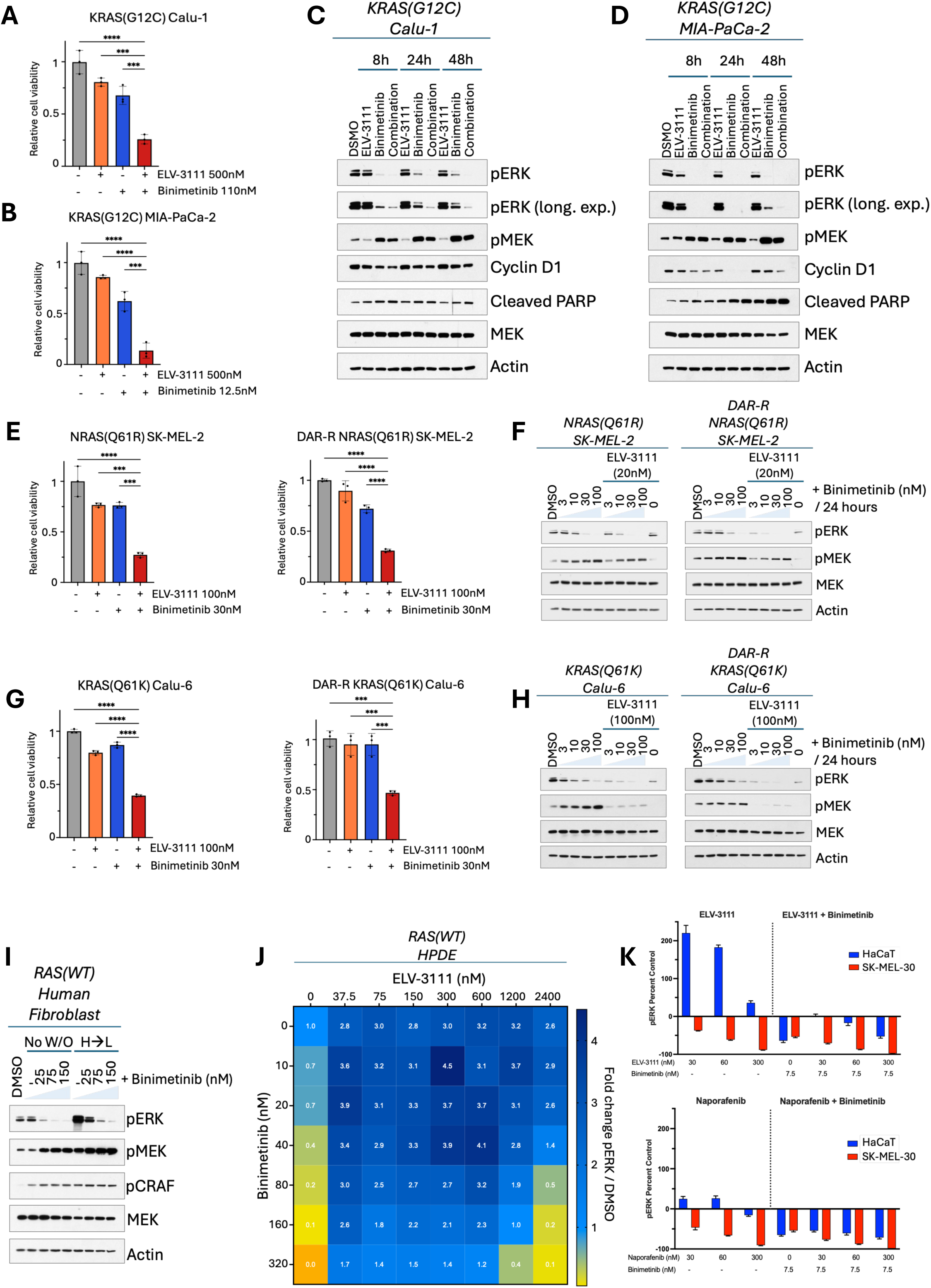
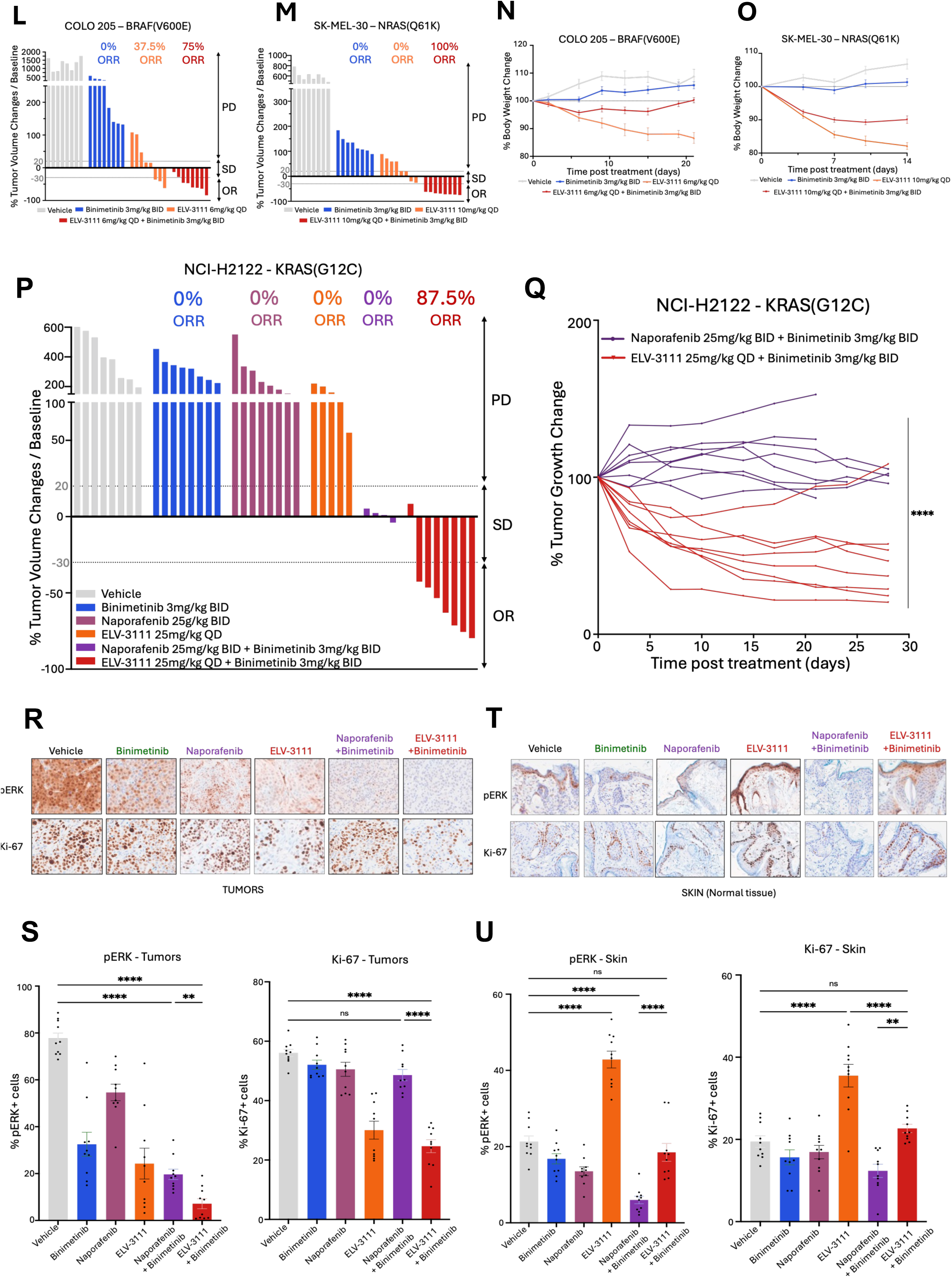
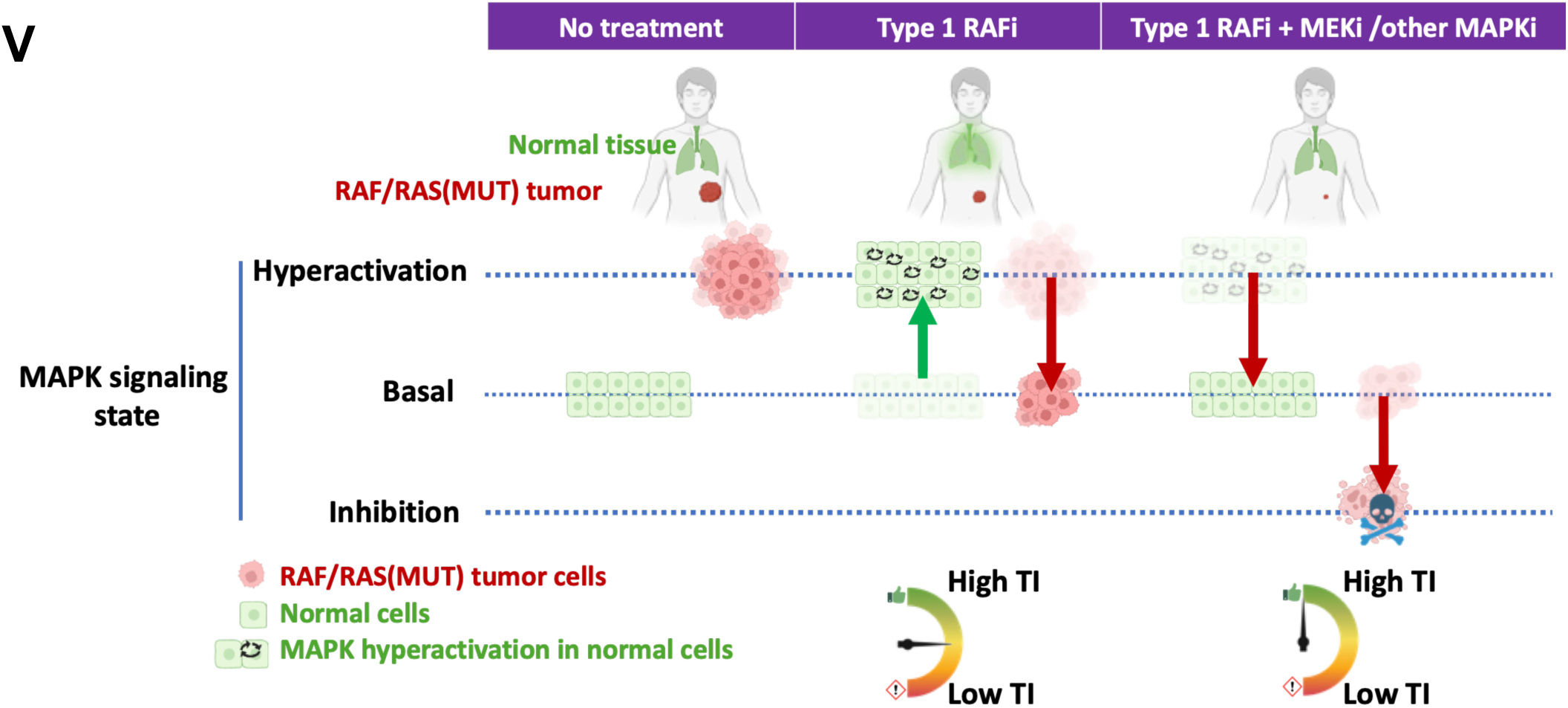
ELV-3111 – based MAPK targeting combinations are highly potent in BRAF(MUT) and RAS(MUT) tumors and well tolerated *in vivo*, including tumor insensitive to current MAPK-targeting therapies. KRAS(G12C) (**A**) Calu-1 and (**B**) MIA-PaCa-2 cell lines were treated for 6 days with 500 nM of Type 1 RAFi ELV-3111, 110nM or 12.5nM of Binimetinib (MEKi) or the combination. Cell viability was measured and represented as relative viability compared to DMSO-treated controls (mean values ± SEM, *n* = 3 independent experiments). KRAS(G12C) (**C**) Calu-1 or (**D**) MIA-PaCa-2 were treated with 500nM of Type 1 RAFi ELV-3111, 100nM of Binimetinib (MEKi) or the combination for 8, 24 or 48 hours. Cell lysates were analyzed by immunoblotting with the indicated antibodies to assess MAPK signaling and cleaved PARP. MEK and actin were used as loading controls. (**E**) Parental NRAS(Q61R) SK-MEL-2 or Daraxonrasib-resistant NRAS(Q61R) SK-MEL-2 (DAR-R SK-MEL-2) cells were treated for 6 days with 100nM of Type 1 RAFi ELV-3111, 30nM of Binimetinib (MEKi), or the combination. Cell viability was measured and represented as relative viability compared to DMSO-treated controls (mean values ± SEM, *n* = 3 independent experiments). (**F**) Parental NRAS(Q61R) SK-MEL-2 or Daraxonrasib-resistant NRAS(Q61R) SK-MEL-2 (DAR-R SK-MEL-2) cells were treated for 24 hours with increasing concentrations of Binimetinib (MEKi) in combination or not with 20nM of Type 1 RAFi ELV-3111. Cell lysates were analyzed by immunoblotting with the indicated antibodies to assess MAPK signaling. MEK and actin were used as loading controls. (**G**) Parental KRAS(Q61K) Calu-6 or Daraxonrasib-resistant KRAS(Q61K) Calu-6 (DAR-R Calu-6) cells were treated for 6 days with 100nM of Type 1 RAFi ELV-3111, 30nM of Binimetinib (MEKi), or the combination. Cell viability was measured and represented as relative viability compared to DMSO-treated controls (mean values ± SEM, *n* = 3 independent experiments). (**H**) Parental KRAS(Q61K) Calu-6 or Daraxonrasib-resistant KRAS(Q61K) Calu-6 (DAR-R Calu-6) cells were treated for 24 hours with increasing concentrations of Binimetinib (MEKi) in combination or not with 100nM of Type 1 RAFi ELV-3111. Cell lysates were analyzed by immunoblotting with the indicated antibodies to assess MAPK signaling. MEK and actin were used as loading controls. (**I**) RAS(WT) Human Fibroblast cells were treated for 8 hours with 500nM (High – H) of ELV-3111 in combination or not with increasing concentrations of the MEKi Binimetinib. After treatment, cells were washed and incubated with 50nM (Low – L) of ELV-3111 and same concentrations of MEKi. MAPK signaling was assessed by immunoblotting. MEK and actin were used as loading controls. (**J**) RAS(WT) HPDE cells were treated with increasing concentrations of Type 1 RAFi ELV-3111 in combination or not with increasing concentrations of Binimetinib (MEKi) for 8 hours. Following treatment, cells were washed and incubated in drug-free conditioned medium for 15 minutes. Cell lysates were analyzed by pERK immunoblotting to assess MAPK signaling. MEK served as a loading control. pERK was quantified, normalized to DMSO control, and represented as a concentration-response heatmap. (**K**) HaCaT (RAF/RAS(WT)) or SK-MEL-30 (NRAS(MUT)) cells were treated with the indicated concentrations of ELV-3111 or Naporafenib with or without Binimetinib for 2 hours, fixed and pERK was measured by in-cell western. pERK percent was normalized to DMSO treated controls (*n* = 3 independent experiments). Mice with established COLO-205 (BRAF(V600E)) or SK-MEL-30 (NRAS(Q61K)) tumors (∼150mm3), were treated p.o. with vehicle, 3 mg/kg Binimetinib BID, 6 mg/kg (COLO-205) or 10 mg/kg (SK-MEL-30) ELV-3111 QD or the combination (*n* = 8 mice per group). Individual tumor volume changes to baseline at the end of the study are represented as waterfall plot for (**L**) COLO-205 and (**M**) SK-MEL-30. Body weight of mice was followed over time and normalized to baseline for (**N**) COLO-205 and (**O**) SK-MEL-30 xenograft-bearing mice. Mice with established NCI-H2122 (KRAS(G12C)) tumors (∼250mm3), were treated p.o. with vehicle, 3 mg/kg Binimetinib BID, 25 mg/kg Naporafenib BID, 25 mg/kg ELV-3111 QD, or Naporafenib + Binimetinib or ELV-3111 + Binimetinib combinations (*n* = 8 mice per group). (**P**) Individual tumor volume changes to baseline at the end of the study are represented as waterfall plot and (**Q**) tumor growth changes for both combinations over time were followed and normalized to the baseline. Mice with established NCI-H2122 (KRAS(G12C)) tumors (∼500mm3), were treated p.o. for 72 hours with vehicle, 3 mg/kg Binimetinib BID, 25 mg/kg Naporafenib BID, 25 mg/kg ELV-3111 QD, or Naporafenib + Binimetinib or ELV-3111 + Binimetinib combinations (*n* = 2 mice per group). Tumor and skin tissues were collected, fixed, and stained for pERK and Ki-67. (**R and T**) Representative IHC staining of tumor and skin sections for pERK and Ki-67. Whole slide images were analyzed for pERK and Ki-67. Five non-overlapping regions of equal area spanning the entire tumor or skin section were quantified to ensure unbiased sampling of the tissue. (**S and U**) Percentage of positive cells for each staining was represented as bar graphs (mean values ± SEM, *n* = 10; ns = not significant (P>0.05), **P<0.01, ****P<0.0001, ordinary one-way ANOVA). (**V**) Therapeutic hypothesis. As a single agent, ELV-3111 demonstrates high potency and selectivity for RAF(MUT) or RAS(MUT) tumors, while induces MAPK hyperactivation in normal cells. This unique pharmacological phenotype creates an opportunity for tumor-selective combination strategies. Combining ELV-3111 with MEK, RAS or SHP2 inhibitors can suppress MAPK hyperactivation in normal tissues, thereby enhancing tumor selectivity and improving the therapeutic index (TI) for RAF/RAS(MUT)-targeted therapies.

The recent advent of direct RAS inhibitors represents a major advance in the treatment of RAS(MUT) cancers, producing meaningful clinical responses across several tumor types(*36, 37*). As with other targeted agents, however, their benefit is frequently limited by acquired resistance and eventual disease progression(*38*). We therefore evaluated the efficacy of the ELV-3111 + Binimetinib combination in NRAS- and KRAS-mutant models with acquired resistance to the multi-RAS(ON) inhibitor Daraxonrasib(*39, 40*). We derived Daraxonrasib-resistant (DAR-R) NRAS(MUT) and KRAS(MUT) cells, which exhibited increased RAS expression and enhanced downstream MAPK signaling. Similar mechanisms of resistance have been reported for mutant-selective RAS inhibitors(*41–43*). These changes conferred resistance to both Daraxonrasib-mediated growth inhibition and MAPK pathway suppression (**fig. S10, H to L**). In contrast, the ELV-3111 + Binimetinib combination suppressed cell growth and MAPK signaling comparably in parental and DAR-R cells (**Fig. 5, E to H**). These results indicate that this combination can be an effective strategy for both treatment-naïve RAS(MUT) and those that have developed resistance to RAS inhibitors, a scenario of increased clinical relevance.

To model the effect of the ELV-3111+MEKi combination on normal cells, we examined MAPK signaling dynamics following treatment and washout in RAS(WT) human fibroblast cells. ELV-3111+MEKi attenuated MAPK hyperactivation after washout, restoring signaling to basal levels (**Fig. 5I**). To define the robustness of MAPK hyperactivation and its buffering using MEK inhibition in normal cells, we performed a checkerboard-style concentration matrix of ELV-3111 and Binimetinib in RAS/RAF(WT) HPDE cells following washout and quantified MAPK dynamics. These data demonstrate that ELV-3111 induces MAPK hyperactivation across a broad concentration range, indicating that this effect is not limited to a narrow exposure window. Importantly, Binimetinib consistently buffers this hyperactivation, maintaining MAPK signaling within an intermediate, physiologically relevant range across clinically relevant concentrations (**Fig. 5J**). Together, these findings define a broad therapeutic buffering window that is robust to variations in drug exposure, supporting the translational feasibility of this combination strategy.

On-target toxicity from MAPK inhibitors in the clinic commonly appears in the skin. We compared the combination of either ELV-3111 or Naporafenib with Binimetinib in a normal human keratinocyte immortalized cell line, HaCaT, and compared it to the NRAS(MUT) melanoma cell line, SK-MEL-30. Binimetinib alone potently suppressed the MAPK pathway in both normal and RAS(MUT) cells (**Fig. 5K**). Naporafenib alone potently inhibited MAPK in the NRAS(MUT) context and modestly suppressed the pathway in the normal cells. However, combined Naporafenib+Binimetinib markedly inhibited MAPK in both contexts, consistent with the dose-limiting toxicities of the Naporafenib+MEKi or other Type 2 RAFi+MEKi combinations observed in the clinic(*17, 44*). ELV-3111 alone inhibited MAPK in the NRAS(MUT) cells but showed MAPK activation in the normal cells. When combined with Binimetinib, ELV-3111 strongly inhibited MAPK in the NRAS(MUT) cells but partially buffered the MAPK suppression in the normal cells (**Fig. 5K**). Thus, the modest MAPK hyperactivation by ELV-3111 without washout *in vitro*, suggests a therapeutic window of the ELV-3111+MEKi compared to Naporafenib+MEKi combination, which can be further expanded with the selective robust MAPK hyperactivation in normal cells *in vivo*.

To assess the effectiveness and tolerability of combinatorial strategies *in vivo*, we tested ELV-3111– based combinations in BRAF(MUT) and RAS(MUT) tumor models. We examined the effect of ELV-3111, alone or in combination with Binimetinib, in mice bearing BRAF(V600E) COLO-205 or NRAS(MUT) SK-MEL-30 xenograft tumors. At a low dose where the free drug concentrations target the *in vitro* pERK IC_50_ in SK-MEL-30 for only 6 hours (0.75mg/kg) (**Fig. 2H**), ELV-3111 alone modestly inhibited tumor growth, while its combination with Binimetinib achieved complete tumor stasis, with modest regressions observed in individual animals, and maintaining good tolerability (**fig. S11, A to C)**. ELV-3111 at higher dose, well above the *in vitro* pERK IC_50_ in SK-MEL-30, or Binimetinib alone promoted either growth inhibition or tumor stasis, respectively. High dose of ELV-3111 together with Binimetinib promoted deep tumor regressions in both BRAF(MUT) and RAS(MUT) mutant contexts, with 75% and 100% of overall response rate (ORR), respectively (**Fig. 5, L and M and fig. S11, D and E**). High doses of ELV-3111 alone were associated with body weight loss, likely a result of MAPK hyperactivation in normal tissues, as it was reversed with the addition of Binimetinib (**Fig. 5, N and O**). To corroborate this tolerability effect with another Type 1 RAFi, we used ELV-1974 (**fig. S6C**). ELV-1974 treatment also induced body weight loss at high concentrations. Similar to ELV-3111 + Binimetinib, the ELV-1974 + Binimetinib combination resulted in deep tumor regressions and completely restored normal body weight (**fig. S11, F and G**). These results show that Type 1 RAFi-mediated MAPK hyperactivation in normal tissues contributes to reduced tolerability *in vivo,* but can be exploited to design safe and effective combinatorial strategies for BRAF/RAS(MUT) tumors.

### ELV-3111, in combination with a MEK inhibitor, induces deep objective responses in a **RAS(MUT) tumor model insensitive to current MAPK-targeting therapies.**

Exclusive reliance on highly responsive preclinical models can substantially overstate the efficacy of RAS/MAPK pathway inhibition, as most RAS(MUT) tumors in both experimental models and clinical settings exhibit, at best, cytostatic responses. This persistent “cytostatic ceiling” reflects a fundamental limitation of current therapeutic strategies, including combined Type 2 RAF and MEK inhibitors, and has posed a major barrier to translating pathway blockade into durable clinical benefit. To determine whether Type 1 RAFi can circumvent this intrinsic resistance and elicit more profound antitumor effects, we employed the NCI-H2122 KRAS(MUT) lung cancer xenograft model, which has consistently demonstrated only growth arrest in response to Type 2 RAFi + MEKi therapy(*15, 45*).

Consistent with previous reports, Naporafenib alone or Naporafenib + Binimetinib combination resulted in progressive disease, or stasis at best (**Fig. 5, P and Q**). ELV-3111 monotherapy likewise resulted in progressive disease, underscoring the necessity of combination treatment. Strikingly, the combination of ELV-3111 with Binimetinib demonstrated markedly superior potency compared to Naporafenib + Binimetinib, reaching an 87.5% Objective Response Rate (ORR), while the Naporafenib + Binimetinib combination elicited no objective responses (**Fig. 5, P and Q**). Although high-dose ELV-3111 monotherapy exhibited reduced tolerability, necessitating dosing holidays and resulting in treatment-related deaths, co-treatment with Binimetinib was well tolerated, with no observed toxicity or excessive body-weight loss. In contrast, the Naporafenib-Binimetinib combination exhibited reduced tolerability, causing rash and necessitating dosing holidays and resulting in animal deaths (4 in the Naporafenib+Binimetinib combinations versus none in the ELV-3111 + Binimetinib treatments) (**fig. S12, A to C**). Reducing the dose of Binimetinib (1.5 mg/kg) or Naporafenib (12.5 mg/kg) resulted in decreased toxicity, as expected; however, it also compromised the antitumor efficacy of the Naporafenib + Binimetinib combination (**fig. S12, D and E**).

In a paired tumor-and-normal-skin pharmacodynamic (PD) analysis performed in the NCI-H2122 xenograft model under the same dosing conditions as the efficacy study, we assessed pERK and proliferative output by immunohistochemistry (IHC) and downstream MAPK signaling by immunoblotting in tumor xenografts and normal skin (**Fig. 5, R to U and fig. S12F**). Across every tumor readout examined, ELV-3111 + Binimetinib matched or exceeded Naporafenib + Binimetinib, while in the same animals it preserved normal-tissue pERK and Ki-67 near baseline, whereas Naporafenib + Binimetinib suppressed them below baseline, and ELV-3111 as a single agent induced MAPK hyperactivation. These PD data link the proposed mechanism, deep tumor suppression with normal-tissue sparing, directly to the drug combination used in the efficacy studies.

Together, these results demonstrate that the combination of ELV-3111 with MEKi induces deep tumor-MAPK-signaling and proliferative output inhibition, while restoring normal MAPK signaling in normal tissues, significantly improving tolerability, which results in markedly increased antitumor efficacy in RAF/ RAS(MUT) tumors (**Fig. 5V**).

## Discussion

Here we uncover a mechanism of MAPK hyperactivation by Type 1 RAFis, selectively in normal cells. Both Type 1 and Type 2 RAFis bind RAF in the αC-helix-IN conformation and thereby drive a shift in the equilibrium towards the active RAF conformation (priming). However, we found that the position of DFG (IN for Type 1 and OUT for Type 2) is associated with markedly different inhibitor dissociation rates from RAF. Thus, drug clearance *in vivo* results in rapid inhibitor dissociation from RAF primed by Type 1 RAF inhibitors, but not Type 2 RAF inhibitors, generating drug-free RAF dimers capable of paradoxically activating MAPK signaling in normal tissues. In contrast, the prolonged residence time of Type 2 RAF inhibitors maintains inhibition of primed RAF despite declining drug concentrations. Importantly, MAPK hyperactivation does not occur in RAS(MUT) tumors because RAF is constitutively primed by oncogenic RAS, rendering it unresponsive to further priming by Type 1 RAF inhibitors. Collectively, these findings identify inhibitor dissociation kinetics, rather than equilibrium target engagement alone, as a key determinant of the biological consequences of RAF inhibition.

A central challenge in targeting the RAS/MAPK pathway has been achieving complete suppression of oncogenic MAPK signaling in tumors while preserving physiological MAPK signaling in normal tissues. This therapeutic ceiling has limited the efficacy of most MAPK-targeted therapies. Current RAF and MEK inhibitor combinations for BRAF(V600X) tumors represent a notable exception. Their success relies on complementary pharmacology, whereby paradoxical MAPK activation induced by Type 1.5 RAF inhibitors is buffered by MEK inhibition in normal tissues while maintaining profound pathway suppression in tumors. The therapeutic opportunity therefore does not derive from paradoxical MAPK activation itself, which represents a dose-limiting liability, but from the ability to buffer this otherwise deleterious effect through vertical MEK inhibition, thereby enabling greater dose intensity and deeper tumor MAPK suppression. Our findings demonstrate that this therapeutic principle is not restricted to Type 1.5 RAF inhibitors or BRAF(V600X) tumors. Instead, the Type 1 RAF inhibitor + MEK inhibitor combination recreates this complementary pharmacology while retaining the ability to inhibit RAF dimers, thereby extending this therapeutic strategy to RAF dimer-driven tumors, including those driven by oncogenic RAS.

Type 1 RAFis were among the first RAF-targeting agents developed, but none advanced to clinical development. This may have reflected insufficient potency or selectivity, as is clearly the case for GDC-0879, rather than an inherent limitation of the Type 1 binding mode itself. ELV-3111 distinguishes itself by exhibiting markedly greater potency and selectivity than earlier Type 1 RAF inhibitors, allowing the unique pharmacology of this class to be therapeutically exploited by combining with a MEK inhibitor. We provide strong preclinical evidence that combining ELV-3111 with a MEK inhibitor induces profound and durable tumor regressions across multiple RAF- and RAS(MUT) models, including a RAS(MUT) model refractory to current MAPK-targeted therapies, while maintaining a favorable tolerability profile. The improved therapeutic index results from complementary pharmacologic effects: enhanced MAPK suppression in tumors together with restoration of physiological MAPK signaling in normal tissues.

Mechanistically, our findings indicate that paradoxical MAPK activation by Type 1 RAF inhibitors is distinct from previously described mechanisms. Multiple alternative explanations, including differential induction of RAF-RAS complexes, RAF dimerization, RAF priming, or ERK-dependent feedback reactivation, were excluded experimentally. Instead, both Type 1 and Type 2 RAF inhibitors similarly promote RAF association with RAS, RAF dimerization, and transition to the primed state, whereas the rapid dissociation of Type 1 RAF inhibitors uniquely permits activation of these primed RAF complexes as drug concentrations decline. This also explains why paradoxical MAPK activation by Type 1 RAF inhibitors is primarily evident *in vivo*, but less apparent under conventional *in vitro* conditions employing constant drug concentrations. Based on its mechanism of action, ELV-3111 provides a versatile backbone for MAPK-targeted combination therapies. Beyond MEK inhibition, combinations with upstream pathway inhibitors, including SHP2(*31*), SOS1(*46*), or RAS(*47*) inhibitors - may be both well tolerated and highly effective, depending on tumor context.

The recent clinical success of direct RAS inhibitors has transformed the treatment of several RAS-mutant cancers. Nevertheless, many RAS-driven tumors remain without effective direct inhibitors, and even in tumors with approved RAS inhibitors, adaptive and acquired resistance commonly restore MAPK signaling through mechanisms involving both mutant and wild-type RAS. In this context, Type 1 RAF + MEK inhibition may provide a complementary therapeutic strategy by targeting RAF downstream of both mutant and wild-type RAS while retaining the therapeutic advantages conferred by selective modulation of MAPK signaling in normal tissues. Consistent with this concept, the ELV-3111 + MEK inhibitor combination remained effective in cellular models with acquired resistance to a multiselective RAS(ON) inhibitor. Although these observations are currently limited to *in vitro* models and require validation *in vivo* and ultimately in clinical studies, they suggest that this strategy may have particular utility in the increasingly relevant post-RAS inhibitor setting, while also providing a therapeutic option for RAS-driven tumors that currently lack effective direct RAS-targeted therapies.

The unique property we discovered may not be exclusive to Type 1 RAFis. Other kinase inhibitors, including those targeting AKT(*48*) or S6K1(*49*), have also been shown to induce target priming. It is conceivable that fast off-rate inhibitors targeting these kinases may similarly promote transient target hyperactivation during drug clearance, creating opportunities for tumor-selective vertical pathway inhibition. More broadly, our findings suggest that signaling dynamics and inhibitor binding kinetics, in addition to target selectivity, can be leveraged to improve the therapeutic index of targeted therapies. Thus, therapeutically exploiting target priming together with rapid inhibitor dissociation through appropriately designed vertical pathway combinations may represent a broadly applicable strategy for overcoming the therapeutic ceiling that has limited therapies targeting oncogenic signaling pathways.

Our study demonstrates that combining a Type 1 RAF inhibitor with MEK inhibition integrates the complementary therapeutic properties of current Type 1.5 and Type 2 RAF inhibitor strategies while broadening the scope of RAF-targeted therapy beyond BRAF(V600X) cancers. By coupling potent inhibition of RAF dimers in tumors with paradoxical MAPK activation selectively in normal tissues, this combination strategy overcomes the therapeutic ceiling imposed by physiological MAPK pathway inhibition, enabling deeper suppression of oncogenic MAPK signaling without a corresponding increase in normal tissue toxicity. Beyond treatment-naïve RAS-driven cancers, this strategy has the potential to expand therapeutic options for tumors that currently lack effective direct RAS inhibitors and for patients whose tumors have progressed following RAS inhibitor therapy, an increasingly common clinical challenge. Together, these findings provide a strong rationale for evaluating Type 1 RAF inhibitor/MEK inhibitor combinations as a broadly applicable therapeutic strategy across RAS/MAPK-driven cancers.

## Acknowledgments

We acknowledge the Cancer Center support grant: NIH/NCI 1P30CA196521-0. We acknowledge Proteros Biosciences for generating the co-crystal of ELV-3111 with BRAF, and Pharmaron for their essential research services in support of this project.

## Funding

P.I.P. is supported by the US National Institutes of Health (NIH) (R01 CA285713, R01 CA240362, R01CA238229), a V Foundation Translational Award, the Melanoma Research Alliance, the Breast Cancer Alliance, the Melanoma Research Foundation, the Irma T. Hirschl Trust and Tisch Cancer Institute developmental awards. M.D is supported by Melanoma Research Foundation (Career Development Award). E.M and E.G. were supported by a National Institutes of Health grant R01CA223243 (E.G), a Montefiore Einstein Comprehensive Cancer Center Support grant P30CA013330, and the Albert Einstein College of Medicine Ph.D. Student Stipend Fund to E.M.

## Author contributions

M.D., T.L.P. conceived and designed the study, acquired, analyzed and interpreted data and drafted and revised the manuscript. E.M., B.B., B.G., A.O.R., L.R, M.R.B, D.D., M.S., acquired, analyzed and interpreted data. J.P.L., S.A.A, contributed to the conception and design of experiments. F.M., E.G. supervised research and interpreted data. P.I.P. and S.D.G. conceived the study, supervised and coordinated research, analyzed and interpreted data and drafted and revised the manuscript.

## Competing interests

P.I.P reports research funding to his institution from Verastem Oncology and Enliven Therapeutics. P.I.P reports consulting fees from Nuvalent Inc, Blueprint Medicines and Belharra Therapeutics. E.G. reports compensation for consulting or equity ownership from BaxGen Therapeutics, BeanPod Biosciences, Comorin Therapeutics, Life Biosciences, Stelexis Biosciences for projects unrelated to this work. T.L.P., L.R., J.R.L., M.R.B., D.D., S.D.G. are employees and shareholders of Enliven Therapeutics. All other authors declare no competing interests.

## Data and Materials availability

All data needed to evaluate the conclusions in the paper are present in the paper or the supplementary materials. The atomic coordinates and structure factors of kinase BRAF (chain A and chain B) bound to ELV-3111 have been deposited in the Protein Data Bank and can be accessed using the accession number 9ZBO.

Supplementary Materials for

## The PDF file includes

Materials and Methods Figs. S1 to S12

Tables S1 to S3

Supplementary Figures and Tables legends References

## Other Supplementary Material for this manuscript includes the following: ELV-3111 Analytical Data

1. LC-MS ELV-3111
2. High-resolution mass spectrometry data for ELV-3111
3. 1H NMR Spectrum of ELV-3111
4. 13C NMR Spectrum of ELV-3111
5. Chiral HPLC analysis of ELV-3111 (100% e.e.)

### PDB validation report

Encorafenib (MedChemexpress, HY-15805), Naporafenib (MedChemexpress, HY-112089), LY3009120 (Selleckchem, S7842), SB-590885 (MedChemexpress, HY-10966), Binimetinib (MedChemexpress, HY-15201), GDC-0879 (MedChemexpress, HY-50864), PLX8394 (Selleckchem, S7965), Belvarafenib (Selleckchem, S8853), Daraxonrasib (MedChemexpress, HY-148439), TNO155 (Selleckchem, S8987), Gefinitib (Medchemexpress, HY-50895), Temuterkib (MedChemexpress, HY-101494) were dissolved in DMSO and stored at −20°C. ELV-3111 and ELV-1974 were developed by Enliven Therapeutics.

### Cloning

For the KRAS-BRAF NanoBiT interaction assay, N-terminal fusions of large BiT (LgBiT) or small BiT (SmBiT) to either full-length KRAS 4A (Uniprot P01116-1) or BRAF (Uniprot P15056) were encoded in pMEV or pCDNA3, respectively. These constructs included 16-residue G/S linkers between the tag and KRAS/BRAF open reading frames (ORFs). The different RAS-mutant used in this study were obtained using QuickChange II XL-site directed mutagenesis kit (Agilent, 200522).

For the NanoBRET assay, BRAF-NanoLuc was obtained using using QuickChange II XL-site directed mutagenesis kit (Agilent, 200522) using BRAF(V600E)-NanoLuc fusion vector (Promega, NV2481).

### Crystallography

BRAF ELV-3111 co-crystallization was performed by Proteros, Inc. Human BRAF kinase domain (amino acids 448-722) with the pFastBac dual vector (ThermoFisher 10712024) and purified by the N-terminal His tag (Cytiva, 17524802) followed by filtration chromatography in 1mM EDTA (Sigma, 798681), 3mM DTT(Sigma, 3860-OP), 15% glycerol (Sigma, G5516) and 25mM Tris HCl pH 8 (Sigma, 648313), with 3mM NaCl (Sigma, S9888), to achieve >95% purity (Cytiva, 28989336). Crystals were generated in 0.1 M HEPES (Thermo, 15630080) pH 7, 0.2M NaCl (Sigma, S9888), 20% PEG 6000 (Sigma, 1546580) buffer by sitting drop method at 1:1 drop ratio at 20 C after several days incubation. X-Ray data was collected at 100K at the Swiss Light Source, beamline PXII, and processed using autoPROC, XDS and autoPROC, AIMLESS^3^. Structure was obtained by molecular replacement, and the model was built and refined using COOT^1^ and CCP4^2^. Crystallographic work was performed by Proteros Biostructures. Structure representations were created using PyMOL (Schrödinger LLC).

### Kinase Screen

A panel of 372 protein and lipid kinases was screened to determine the percent inhibition of these kinases employing 1 mM tested compound in duplicate with 100 mM ATP (Reaction Biology). Kinases that were inhibited by >50% were considered hits and therefore selected for follow-on IC50 determinations.

### Cell culture

*Cell lines*

All cell lines were cultured under standard conditions (37 °C, 5% CO₂).

SK-MEL-239, SK-MEL-239 C3, MEFs, HeLa, primary human fibroblasts, A375, COLO 205, COLO 201, HEK293, COS-7, AsPC-1, MiaPaCa-2 were maintained in DMEM (ThermoFisher, 10313-039) supplemented with 10% fetal bovine serum (Biowest, S1680), 2 mM glutamine (ThermoFisher, 25-030-164) and 100U/mL penicillin/streptomycin (ThermoFisher, 15-140-163).

SK-MEL-264, SK-Mel-246, SK-MEL-2, Calu-1, NCI-H358, SK-MEL-30, WM1789, WM3928, WM3670, WM3912, WM3629, WM3406, WM3844, HMVI, Calu-6 and Ba/F3 were maintained in RPMI (ThermoFisher, 11-875-119) with the same supplements.

HPAC cells were maintained in DMEM/F12 (ThermoFisher, 11320033) with the same supplements. OCI-AML-2 were cultured in Mem Alpha (Gibco, 12571-063) with the same supplements. Caco-2 were cultured in EMEM (Sigma, M4655) with the same supplements. HPDE were cultured in Keratinocyte SFM, + EGF + bovine pituitary extract (Invitrogen, 17005042). Hermes 2B were cultured in Ham’s F-10 medium (ThermoFischer, 11550043), supplemented with 10% FBS, 2 mM L-glutamine, 200 pM cholera toxin (Sigma-Aldrich, C8052), 10 ng/mL human SCF (Life Technologies, PHC211), 10 nM endothelin-1 (Sigma-Aldrich, E7764), and 200 nM TPA (Sigma-Aldrich, P8139).

### Transient Transfection

HEK293 cells were seeded in 60mm dishes to reach 70-80% confluency at the time of transfection. For each dish, 1μg of plasmid DNA was diluted in 250 μL of serum-free DMEM. Fugene HD transfection reagent (Promega, E2311) was added at a 1:3 DNA-to-reagent ratio. The mixture was gently pipetted to mix and incubated at room temperature for 10–15 minutes to allow complex formation. The transfection mixture was added dropwise to the cells. Cells were maintained under standard culture conditions (37 °C, 5% CO₂) and harvested 24-48 hours post-transfection for downstream assays.

### Lentivirus production and Cell Transduction and doxycycline-inducible stable cell line generation

Lentiviral particles were generated by transfecting HEK293T cells with the pTRI-BlaS transfer plasmid encoding the KRAS(Q61K) variant, along with the packaging vector psPAX2 (Addgene), and envelope vector pMD2.G (Addgene) at a 5:4:1 ratio, using Lipofectamine 2000 (ThermoFisher, 11668019). The viral supernatant was collected 72 hours post-transfection, filtered through a 0.45 μm filter unit (MilliporeSigma, SLHPR33RS), and used to infect HeLa and MEFs in the presence of 10 μg/mL polybrene (Sigma-Aldrich, TR-1003). Infected cells were incubated overnight at 37°C, after which viral supernatant was removed and replaced with fresh medium. Transduced cells were selected by culturing in 3 μg/ml Blasticidin (Invivogen, ant-bl-05) for approximately 7 days.

Prior to use in experiments, HeLa and MEFs cells were treated with 500 ng/ml or 1000 ng/mL doxycycline, respectively, for 48 hours.

### COLO 201 Resistant cells

The COLO 201 cell line was engineered to homozygously express either NRAS(Q61K) or KRAS(G12V) using CRISPR/Cas9. The design and synthesis of CRISPR components, transfection, single clone isolation and confirmation was done at Synthego.

### Ba/F3 Screen

Engineered Ba/F3 cell lines expressing either BRAF Class I (V600E), Class IIa (K601N, L597R), Class IIb (G469A, G464R), or Class IIc (MAD1L1-BRAF, AGAP3-BRAF fusions) mutations were tested in 72-hour proliferation assays using a dose response of ELV-3111 (Kyinno Biotechnology).

### Cellular viability assay

For the initial characterization of ELV-3111 (Fig. 1C), cells were plated at a density of 0.5-1.5.10^3^ cells/well in a 384 well plate (Corning™, 3570) or 3000 cells/wells in 96-well plates (Corning™, 3904) and allowed to adhere overnight. The following day, cells were treated with a dose response of the indicated compounds and incubated for 72-120h, depending on the cell line. CellTiterGlo (Promega G9241), was added per manufacturer’s protocol and viability was calculated relative to DMSO-treated controls and GI50 values were calculated using non-linear regression 4-parameter fit algorithm in Graphpad Prism.

For all other cell viability assessments, cells were seeded in 96-well plates (Corning™, 3595) at a density of 0.5-1.10³ cells per well. The following day, cells were treated with the indicated inhibitors in standard growth medium. After six days of treatment, 10 µL of Cell Counting Kit-8 solution (Dojindo Molecular Technologies, CK04) was added to each well, and plates were incubated at 37 °C for 2.5 hours. Absorbance was measured at 450 nm using a plate reader (Promega GloMax® Discover Microplate Reader, GM3000). Dose-response curves and GI50 values were generated using log-transformed and normalized data.

For combination experiments, compounds were added in a two-dimensional dose response. Relative viability was uploaded into SynergyFinder 3.0^4^ and scored using the most synergistic area method.

### Western blot and active RAS pull-down

Immunoblotting was performed using standard methods. Cells were washed with ice-cold PBS and lysed on ice in NP-40 lysis buffer (50 mmol/L Tris pH 7.5, 1% NP40, 150 mmol/L NaCl, 10% Glycerol 1 mmol/L EDTA) supplemented with protease inhibitors (Roche, 11873580001) and phosphatase inhibitors (Roche, 4906837001). Lysates were centrifuged at 15,000 rpm for 15 minutes at 4°C, and the protein concentrations were determined by BCA (ThermoFisher, 23221, 23224). Equal amounts of proteins were resolved by SDS-PAGE on NuPAGE 4-12% Bis-Tris Gels (Invitrogen, NP0321BOX), and transferred to nitrocellulose membranes (GE Healthcare, 45-004-006). After blocking in 5% Milk, membranes were immunoblotted overnight with antibodies against: pMEK1/2Ser217/22 (Cell Signaling, 9154), MEK1 (Cell Signaling, 2352), pERK1/2Thr202/Tyr204 (Cell Signaling, 4370), pCRAF (Millipore – Sigma, 05-538), pEGFR (Cell signaling, 2234), pRb (Cell signaling, 9307), CRAF (BD Transduction Laboratories, 610152), BRAF (Santa-Cruz, sc-5284), RAS (ThermoFisher Scientific, 1862335), NRAS (Santa Cruz, sc-31), KRAS (Proteintech, 12063-1-AP), cleaved-PARP (Cell Signaling, 9541), DUSP4 (Cell signaling, 5149), DUSP6 (Cell signaling, 50945), Cyclin D1 (Cell Signaling, 2922). The next day, membranes were probed with anti-rabbit IgG (Cell Signaling, 7074) or anti-mouse IgG (Cell Signaling, 7076) secondary antibodies, and chemiluminescent signals were detected on X-ray films. Loading controls were revealed using b-actin HRP-conjugate (Cell Signaling, 5125S).

RAS pull-down kit (ThermoFisher, 16117) was used to determine levels of RAS-GTP, according to the manufacturer’s protocol. RAS was detected by western blot using pan-RAS antibody provided with the kit.

### pERK In-Cell Western

Cells were plated to a density of 70-90% in 96 well plates (Corning 3904) and allowed to adhere overnight. The following day, cells were treated with a dose response of the indicated compounds for two hours and then fixed with 4% paraformaldehyde (FisherScientific BP531) for 20 minutes. After fixation, cells were washed with PBS and permeabilized with ice-cold methanol for 10 minutes. Non-specific binding was blocked with blocking buffer (Licor 927-60001) before overnight incubation with primary antibodies pERK1/2Thr202/Tyr204 (Cell Signaling, 9101) and GAPDH (Millipore MAB74) at 4°C. Secondary antibodies or CellTag (Licor 700/800 anti-Rabbit/Mouse or 926-41090) were incubated for 1 hour at room temperature. Fluorescent signal was detected on the Odyssey M (Licor) and pERK signal was normalized to GAPDH/CellTag for each well. pERK was calculated relative to DMSO-treated controls and IC50 values were calculated using non-linear regression 4-parameter fit algorithm in Graphpad Prism.

For human serum shift quantification normal cell media was replaced with 100% human serum (BioVT HUMANSRM-0001255) immediately prior to treatment.

### TR-FRET Phospho-ERK1/2

Phospho-ERK1/2 (T202/Y204) levels were measured using the THUNDER™ Phospho-ERK1/2 TR-FRET Cell Signaling Assay Kit (Bioauxilium, KIT-ERKP-500 / KIT-ERKT-500) according to the manufacturer’s instructions.

### Co-immunoprecipitations

500–1000μg of proteins from cell lysates were pre-cleared with Protein G agarose (ThermoFisher, 15-920-010) for 2 hours at 4°C. Pre-cleared lysates were incubated with 2μg BRAF antibody (Santa-Cruz, sc-5284) and Protein G agarose overnight at 4°C. Samples were then washed three times with lysis buffer and sample buffer was added for subsequent immunoblot analysis.

### Washout experiments

For complete washout experiments, cells were seeded in 60mm dishes at a density of 1.10^6^ cells per plate. The following day, cells were treated for 8 hours with saturating concentrations of RAFis (1μM for all compounds, except GDC-0879 – 10μM). After treatment, cells were washed with pre-warmed 37°C 1X-PBS, and drug-free conditioned growth medium was added. Cells were then incubated for the indicated durations post-washout and analyzed by western blot.

For washout experiments based on PK/PD studies, cells were seeded in 60mm dishes at a density of 1.10^6^ cells per plate. The following day, cells were treated for 8 hours with 500 nM or 150 nM of RAFis ELV-3111 or LY3009120. After treatment, cells were washed with pre-warmed 37°C 1X-PBS and treated with 50 nM or 4 nM of compounds, respectively. Cells were then incubated for 30 minutes and analyzed by western blot.

### In-cell Nano-BiT

HEK293 or COS 7 cells were co-transfected with constructs using FuGENE HD (Promega, E2311) and 10,000 cells were cultured overnight in white 96-well plates (ThermoFisher, 236105) at 37°C. Plates were then treated with different RAFis and 20 μM fluorofurizamine. Relative luminescence was determined using a plate reader (Promega GloMax® Discover Microplate Reader, GM3000).

### In vitro Binding kinetic profiling

A 96-well plate was prepared using serial dilutions of each compound in 100% DMSO at 100x the final test concentration. From these, final assay concentrations of 10 nM and 100 nM were selected for analysis. Wells containing 100% DMSO were used as high and low controls.

For assay preparation, 1.5 µL of each compound dilution was transferred into a dilution plate containing 48.5 µL of assay buffer (50 mM HEPES, pH 7.5; 1 mM EGTA; 0.01% Brij-35; 10 mM MgCl₂; 2 mM DTT). To minimize precipitation, dilutions were prepared immediately prior to initiating the assay. Subsequently, 5 µL of each solution was transferred into the assay plate to reach the desired final concentrations (10 or 100 nM) in 1% DMSO. Test compounds and the fluorescent probe were added simultaneously to a final reaction volume of 15 µL, and fluorescence was recorded for 240 minutes at room temperature. Each compound was tested in singlicate. Binding kinetics were measured using the KINETICfinder® TR-FRET Binding Kinetic Assay (Enzymlogic).

### NanoBRET experiments

*Kinetic analysis of target engagement via BRET*

2.10^6^ HEK293 cells were transfected in suspension with 4 μg of BRAF-NanoLuc plasmid DNA and either 1 μg of NRAS(Q61K) plasmid or 1 μg of Transfection Carrier DNA (Promega, E4881), using 15 μL of FuGENE HD (Promega, E2311). Transfected cells were plated in 96-well white plates (Thermo Fisher, 236105) at a density of 2 × 10^4^ cells per well and incubated overnight at 37 °C. Next day, cells were treated with 62.5 nM LY-Tracer and increasing concentrations of tested RAFis for 2 hours. 20 μM fluorofurizamine was added to each well and the relative BRET signal was determined using a plate reader (Promega GloMax® Discover Microplate Reader, GM3000).

To compare RAF isoform selectivity, the K4 tracer was used with BRAF or CRAF. For the ARAF assay a BRAF/ARAF heterodimer was expressed and displacement of the K10 tracer was measured (Promega).

### Kinetic analysis of ligand dissociation via BRET

2.10^6^ COS-7 cells were transfected in suspension with 4 μg of BRAF-NanoLuc plasmid DNA and 1 μg of Transfection Carrier DNA (Promega, E4881), using 15 μL of Fugene HD (Promega, E2311) transfection reagent. Transfected cells were plated in white 96-well plates (Thermo Fisher, 236105) at a density of 2 × 10^4^ cells per well and incubated overnight at 37 °C. Next day, cells were treated with saturating concentrations of tested RAFis (∼10-fold BRET ratio IC50) for 2 hours. After treatment, cells were washed or left unwashed with pre-warmed 37°C 1X-PBS, 62.5 nM of LY-tracer diluted in conditioned growth medium and 20 μM of fluorofurimazine was added into each well. Relative BRET signal was determined over time using a plate reader (Promega GloMax® Discover Microplate Reader, GM3000).

### RAF protomer dimerization BRET

RAF dimerization was assessed using the NanoLuc® and HaloTag® Bioluminescence Resonance Energy Transfer (BRET) system (Promega). HEK293 cells were resuspended at a density of 2 × 10⁵ cells/mL and transiently transfected using FuGENE® transfection reagent according to the manufacturer’s instructions. For isoform selectivity experiments, cells were transfected with a plasmid ratio of 9:1 (KRAS G12C:RAF expression constructs). For general dimerization experiments, carrier plasmid was substituted for KRAS G12C.

In BRAF/CRAF heterodimer assays, BRAF was fused to NanoLuc® and CRAF to HaloTag®. BRET measurements were performed using the NanoBRET™ Nano-Glo® Substrate and the HaloTag® NanoBRET™ 618 Ligand (Promega N2011 and G9801). Signal detection was carried out according to the manufacturer’s protocol, and BRET efficiency was quantified as a measure of RAF protomer dimerization.

### Caspase-Glo 3/7 assay

Cells were seeded at a density of 10,000 cells/well in 96-well white plates. After 24 hours, drug treatment or vehicle was added, and cells were assayed as indicated. Apoptosis was measured using the Promega Caspase-Glo 3/7 Assay System (G8090) as per the manufacturer’s instructions.

### Immunohistochemistry

For immunohistochemical analysis, tumor xenografts and normal skin tissue were harvested and fixed in 10% aqueous buffered zinc formalin and paraffin embedded. Tissue processing and staining were performed as described previously using the following antibodies: Ki67 (Cell Signaling Technologies, 12202), pERK (Cell Signaling Technologies, 4370). Samples were mounted in prolong gold anti-fade mounting medium (Invitrogen) and were scanned using a Zeiss Axioscan Z1 slide scanner equipped with a 20×/0.8 NA objective. All image analyses were performed using the QuPath software to perform cell detection and quantify each marker as a percentage of stained cells.

### Molecular Dynamics Simulations

Molecular dynamics (MD) simulations were carried out using DESMOND (Schrödinger, LLC, 2023–24). The crystal structures of ELV-311, LY3009120 (PDB ID 5C9C), GDC-0879 (PDB ID: 4MNF) and Sorafenib (PDB ID: 1UWJ) in complex with BRAF were prepared using the Desmond System Builder. Each complex was solvated in a TIP3P explicit water model using a truncated orthorhombic box with periodic boundary conditions. For ELV-3111, the system contained 34,760 atoms including 10,066 water molecules and was neutralized with 28 Na⁺ and 35 Cl⁻ counterions to achieve a final ionic strength of 0.15 M NaCl. For LY3009120, the system contained 30,657 atoms including 8,716 water molecules and was neutralized with 24 Na⁺ and 33 Cl⁻ counterions at 0.15 M NaCl. For GDC-0879, the system contained 32,357 atoms including 9,287 water molecules and was neutralized with 26 Na⁺ and 35 Cl⁻ counterions to achieve a final ionic strength of 0.15 M NaCl. For Sorafenib, the system contained 33,066 atoms including 9,499 water molecules and was neutralized with 26 Na⁺ and 34 Cl⁻ counterions at 0.15 M NaCl. Ligand parameters were assigned with the OPLS4 force field (ELV-3111: C₂₂H₂₁FN₆O₂, 420.45 Da, 52 atoms; LY3009120: C₂₃H₂₉FN₆O, 424.53 Da, 60 atoms, GDC-0879: C19H18N4O2, 334.38 Da, 43 atoms;

Sorafenib: C21H17ClF4N4O3, 484.84 Da, 50 atoms).All systems were subjected to energy minimization and relaxation protocols prior to production. MD simulations were performed under NPT ensemble conditions at 300 K and 1.0325 bar, with temperature and pressure maintained using the Nose–Hoover chain thermostat and Martyna–Tobias–Klein barostat, respectively. Each simulation was run for 1,000 ns, with trajectories recorded every 250 ps. Three independent simulations with different random seeds were performed for ELV-3111, LY3009120, GDC-0879, and Sorafenib. Trajectories were analyzed using Desmond Simulation Interactions Diagram tools (Schrödinger, LLC). Root mean square deviation (RMSD), root mean square fluctuation (RMSF), secondary structure evolution, and protein–ligand contact frequencies were calculated over the course of the simulations. Structural analyses and overlays were performed using Maestro and PyMOL (Schrödinger, LLC).

### In vivo Efficacy Studies

SK-MEL-30 and COLO-205 cell lines were cultured in RPMI 1640 medium supplemented with 10% fetal bovine serum (FBS) at 37°C in a humidified incubator with 5% CO₂. SK-MEL-30 cells were grown as a monolayer, while COLO-205 cells were maintained as a mixed monolayer suspension. Cells in exponential growth phase were harvested, counted, and resuspended for inoculation. Female Balb/c nude mice were subcutaneously inoculated in the right flank with either 1 × 10⁷ SK-MEL-30 cells in 0.1 mL of a 1:1 RPMI 1640:Matrigel mixture, or 5 × 10⁶ COLO 205 cells in 0.1 mL RPMI 1640.

Compounds were formulated in 0.5% methylcellulose/0.5% Tween-80 in water and administered by oral gavage. Mice were monitored biweekly for tumor growth using digital calipers and observed regularly for general health indicators, including body weight, grooming behavior, mobility, and food and water intake.

NCI-H2122 cell line was cultured as monolayer in RPMI 1640 medium supplemented with 10% fetal bovine serum (FBS) at 37°C in a humidified incubator with 5% CO₂. Cells in exponential growth phase were harvested, counted, and resuspended in PBS for inoculation. Female athymic Nude-Foxn1nu mice were subcutaneously inoculated in the right flank with 1 × 10⁷ cells. All mouse experiments were approved by the Institutional Animal Care and Use Committee (IACUC) of WuXi AppTec, Shanghai ChemPartner or the Icahn School of Medicine at Mount Sinai (protocol no. IACUC–2016–0066). Mice were maintained under specific pathogen-free conditions, and food and water were provided ad libitum. ELV-3111 and Binimetinib were resuspended in 0.5% methylcellulose/0.5% Tween-80 in water, Naporafenib compound was resuspended in 40%PEG400 / 10% ethanol in water, and administered by oral gavage. Mice were monitored biweekly for tumor growth using digital calipers and observed regularly for general health indicators, including body weight, grooming behavior, mobility, and food and water intake.

### Pharmacokinetic studies in Balb/c nude mice

In brief, female Balb/c nude mice were administered the indicated dose of ELV-3111 via oral gavage in 0.5% methylcellulose/0.5% Tween-80 in water at 5 mL/kg. Blood samples were collected at various time points from 2-24h after administration for processing to plasma and subsequent analysis by LC/MS/MS to determine compound concentration.

### Pharmacodynamic study in Tumor-Bearing Mice

SK-MEL-30 tumors were established in Balb/c nude mice as described above. Once tumors reached approximately 300 mm³, mice received a single oral dose of test compounds, either alone or in combination. ELV-3111 was formulated in 5% TPGS, Binimetinib in 0.5% methylcellulose/0.5% Tween-80, and Naporafenib in 40% PEG 400/10% ethanol/50% water.

Tumors, kidneys, and lungs were harvested post-treatment and flash-frozen for downstream analyses. Tissues were homogenized in RIPA buffer using a bead-based tissue homogenizer and centrifuged at 10,000 × g for 10 minutes at 4°C. Supernatants were adjusted to a final protein concentration of 4 μg/μL with RIPA buffer and prepared for SDS-PAGE by the addition of 1× Laemmli sample buffer.

Proteins were separated by SDS-PAGE, transferred to nitrocellulose membranes, and probed with antibodies specific for total and phosphorylated ERK. Detection was performed using infrared fluorophore-conjugated secondary antibodies and visualized using a LI-COR Odyssey imaging system. Signal intensities were quantified to assess target modulation.

### Quantification and Statistics

Western blot quantifications were performed using ImageJ (NIH). Intensity values were normalized to loading controls and expressed relative to control conditions. Data were analyzed using GraphPad Prism. Statistical analyses and tests are detailed in the figure legends.

---

**Fig. S1.** ELV-3111 is a Type 1 RAF inhibitor with broad activity and potent inhibition of both monomeric and dimeric RAF. (**A–F**) 2mFo–DFc maps contoured at 0.7 rmsd are shown as gray mesh, illustrating continuous density supporting the ligand model. mFo–DFc difference maps, contoured at ±3 rmsd, are displayed in green (positive) and purple (negative). The ligand is shown in multiple orientations to visualize fit and density quality. (**G**) Overall structure of the BRAF kinase domain in complex with ELV-3111, highlighting the ligand position within the ATP-binding cleft. (**H**) Close-up view of the ELV-3111 binding pocket. The 2mFo–DFc map (blue mesh) confirms the accuracy of ligand placement. Electron-density maps were generated using Phenix and visualized in PyMOL.

**Fig. S2.**
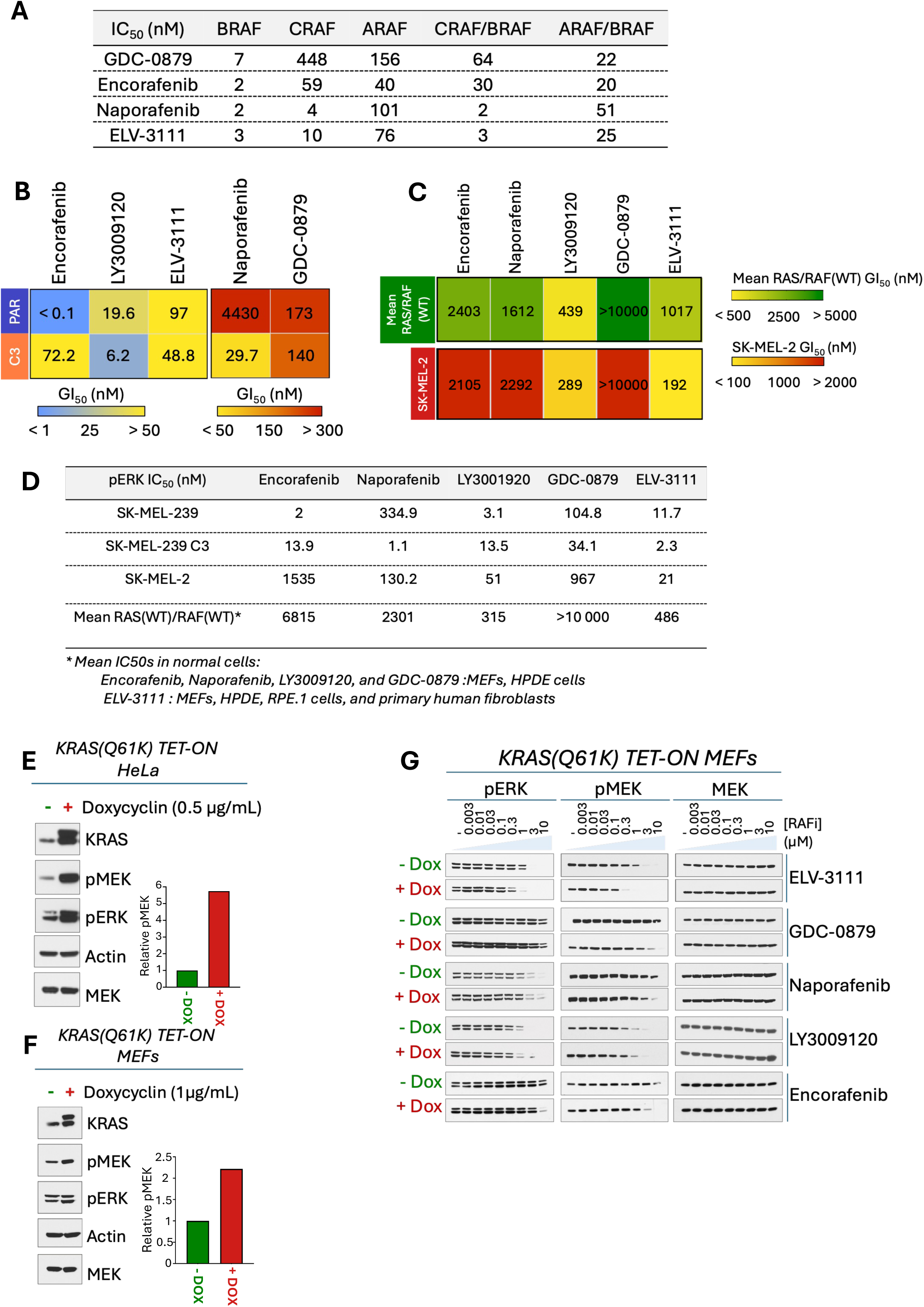
ELV-3111 is a Type 1 RAF inhibitor with broad activity and potent inhibition of both monomeric and dimeric RAF. (**A**) NanoLuc tagged A/B/CRAF was expressed in HEK293 cells and treated with K10 (ARAF) or K4 (B/CRAF) tracer and a dose range of the indicated compounds for 2 hours. IC_50_s were generated from BRET measurements (geometric mean, *n* = 3 independent experiments). Ratios of the indicated IC_50_s are shown in the last two columns. GI_50_s of cell viability after treatments of either (**B**) parental SK-MEL-239 or clone C3 SK-MEL-239, (**C**) Mean of RAF/RAS(WT), tested in Fig. 1F, or SK-MEL-2 cells are represented as heatmaps with corresponding values representing means of 3 independent experiments. (**D**) pERK IC_50_s of response of the indicated cell lines to RAFis was calculated from the ratio of phosphorylated ERK to MEK and represented as a table. (**E**) Doxycycline-inducible KRAS(Q61K) _HeLa_ cells were treated with doxycycline (500ng/mL) for 48 hours. Samples were analyzed by immunoblotting with the indicated antibodies to assess KRAS expression and MAPK signaling. MEK and actin were used as loading controls. (**F**) Doxycycline-inducible KRAS(Q61K) MEFs cells were treated with doxycycline (1μg/mL) for 48 hours. Samples were analyzed by immunoblotting with the indicated antibodies to assess KRAS expression and MAPK signaling. MEK and actin were used as loading controls. (**G**) Doxycycline-inducible KRAS variant MEFs cells were treated with doxycycline (1μg/mL) for 48 hours to induce KRAS(Q61K) expression, followed by 1-hour treatment with increasing concentrations of different types of RAFis. Cell lysates were immunoblotted with the indicated antibodies to assess MAPK signaling. MEK was used as loading control. Exposure times were adjusted separately for −Dox and +Dox conditions to visualize treatment effects; baseline signaling is shown in fig. S2, E and F.

**Fig. S3.**
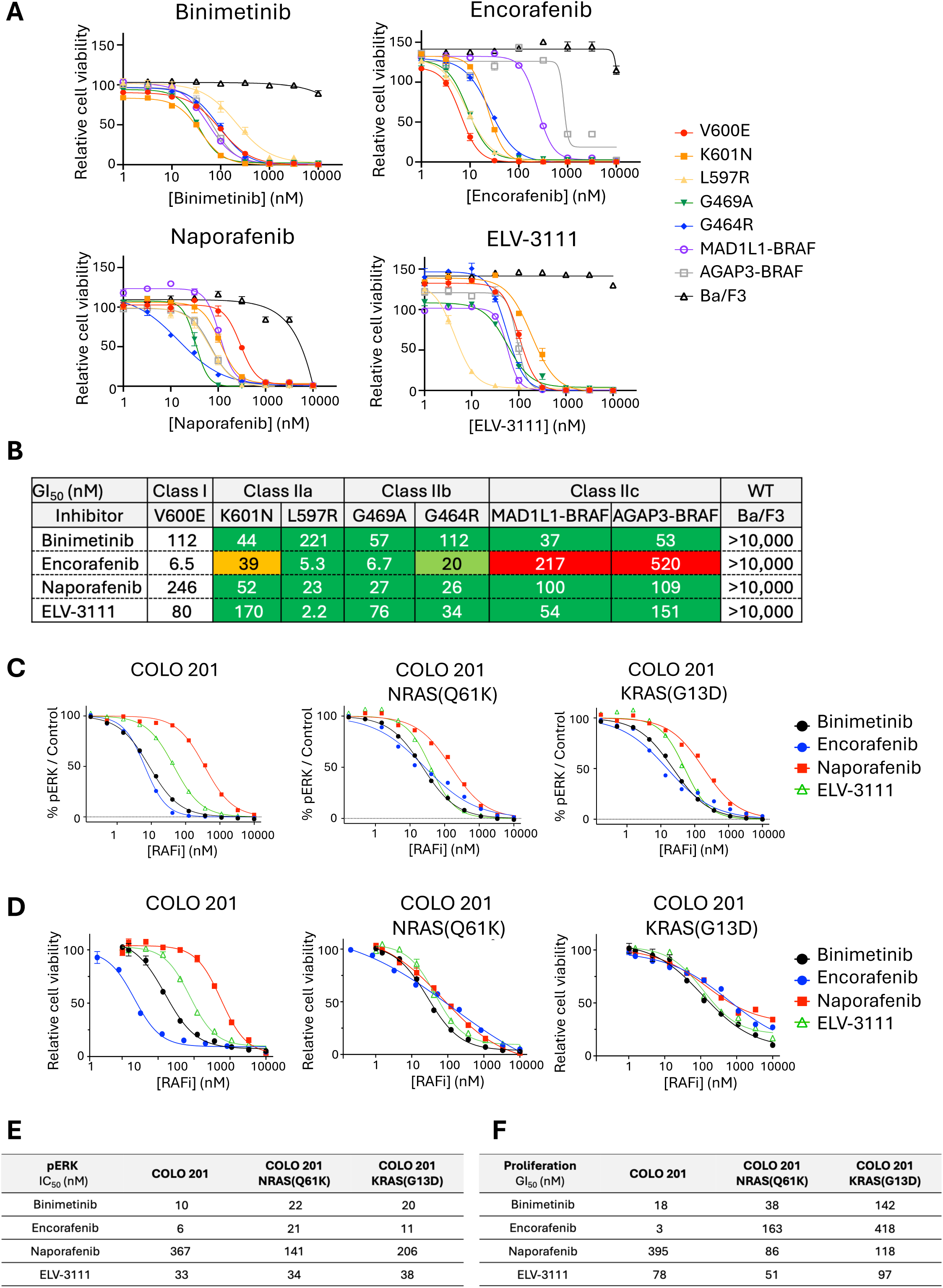
ELV-3111 is a Type 1 RAF inhibitor with broad activity and potent inhibition of both monomeric and dimeric RAF. Ba/F3 cell lines expressing the indicated BRAF mutants were treated with a range of inhibitor concentrations of the indicated RAF or MEK inhibitors for 72 hours. (**A**) Viability was measured and represented as relative cell viability compared to DMSO-treated controls (*n* = 3 independent experiments). (**B**) GI_50_s are represented as table. (*n* = 3 independent experiments). BRAF(V600E) COLO 201 cells were engineered to express NRAS(Q61K) or KRAS(G13D) mutations. Profiling data for ELV-3111 and other inhibitors were calculated from dose responses. (**C**) An in-cell western procedure was used to measure pERK inhibition after 2 hours of treatment. (**D**) Proliferation assays were performed in the indicated cell lines after 5 days of treatment. (**E**) pERK IC_50_s or (**F**) GI_50_s were represented as a table (geometric mean, *n* = 3 independent experiments).

**Fig. S4.**
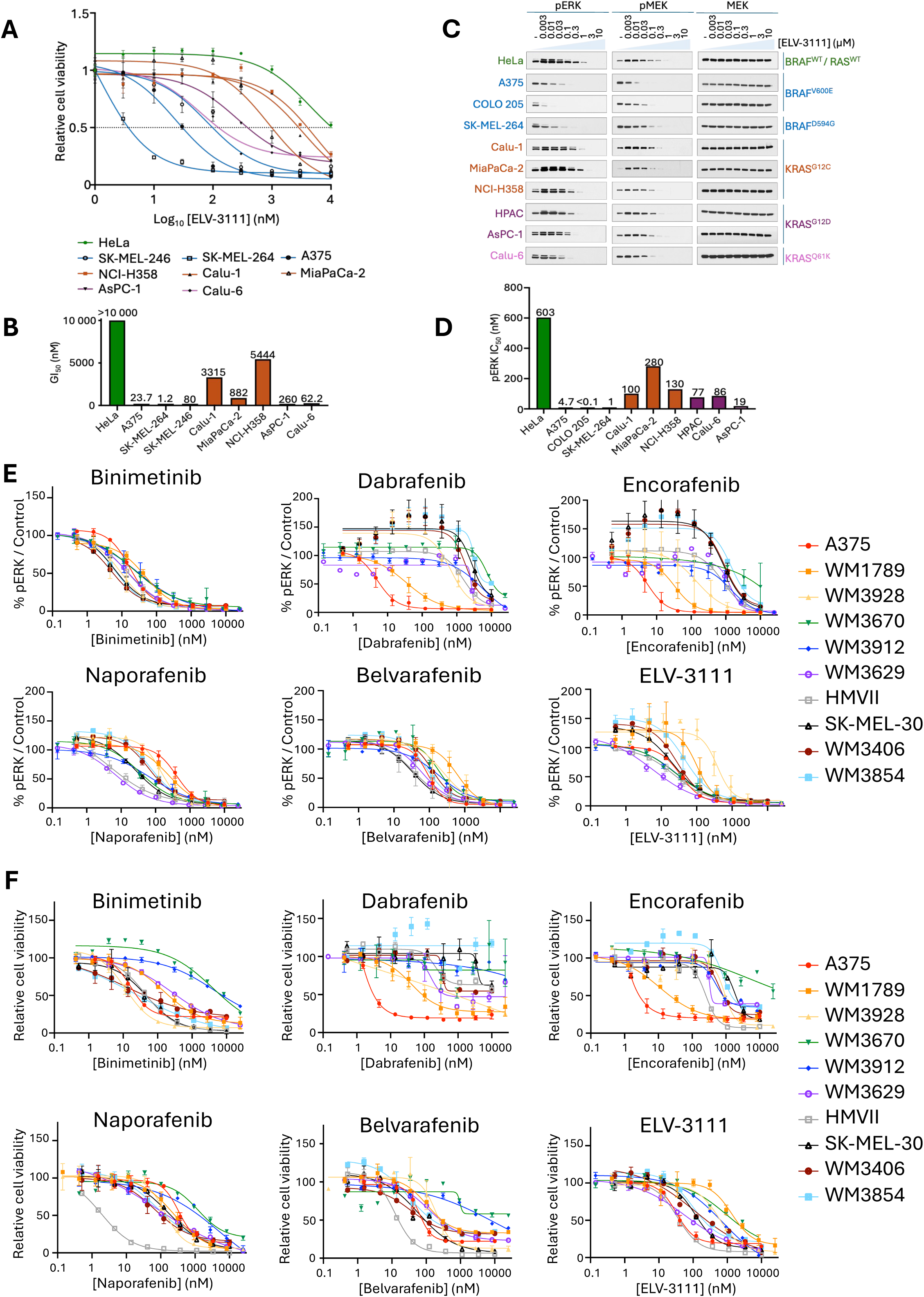
ELV-3111 is a Type 1 RAF inhibitor with broad activity and potent inhibition of both monomeric and dimeric RAF. A panel of cell lines expressing either RAS(WT), different classes of BRAF mutations or RAS mutations was treated for 6 days with increasing concentrations of ELV-3111. (**A**) Cell viability was measured after 6 days of treatment and represented as relative viability compared to DMSO-treated controls (mean values ± SEM, *n* = 3 replicates). (**B**) GI_50_s after treatments were represented as bar graphs. (**C**) Effect on MAPK signaling was assessed by immunoblotting after 1 hour of treatment with ELV-3111. MEK were used as loading control. (**D**) Ratio of phosphorylated ERK to MEK was calculated and pERK IC_50_s after treatment are represented as bar graphs. Cell lines with a spectrum of BRAF and NRAS mutations were screened for sensitivity to ELV-3111 and other MAPK pathway inhibitors. (**E**) An in-cell western procedure was used to measure pERK inhibition after 2 hours of treatment with a dose range of different RAFis. (**F**) Proliferation assays were performed in the indicated cell lines after 72h (A375) or 120h (all other lines) (geometric mean, *n* = 3 independent experiments).

**Fig. S5.**
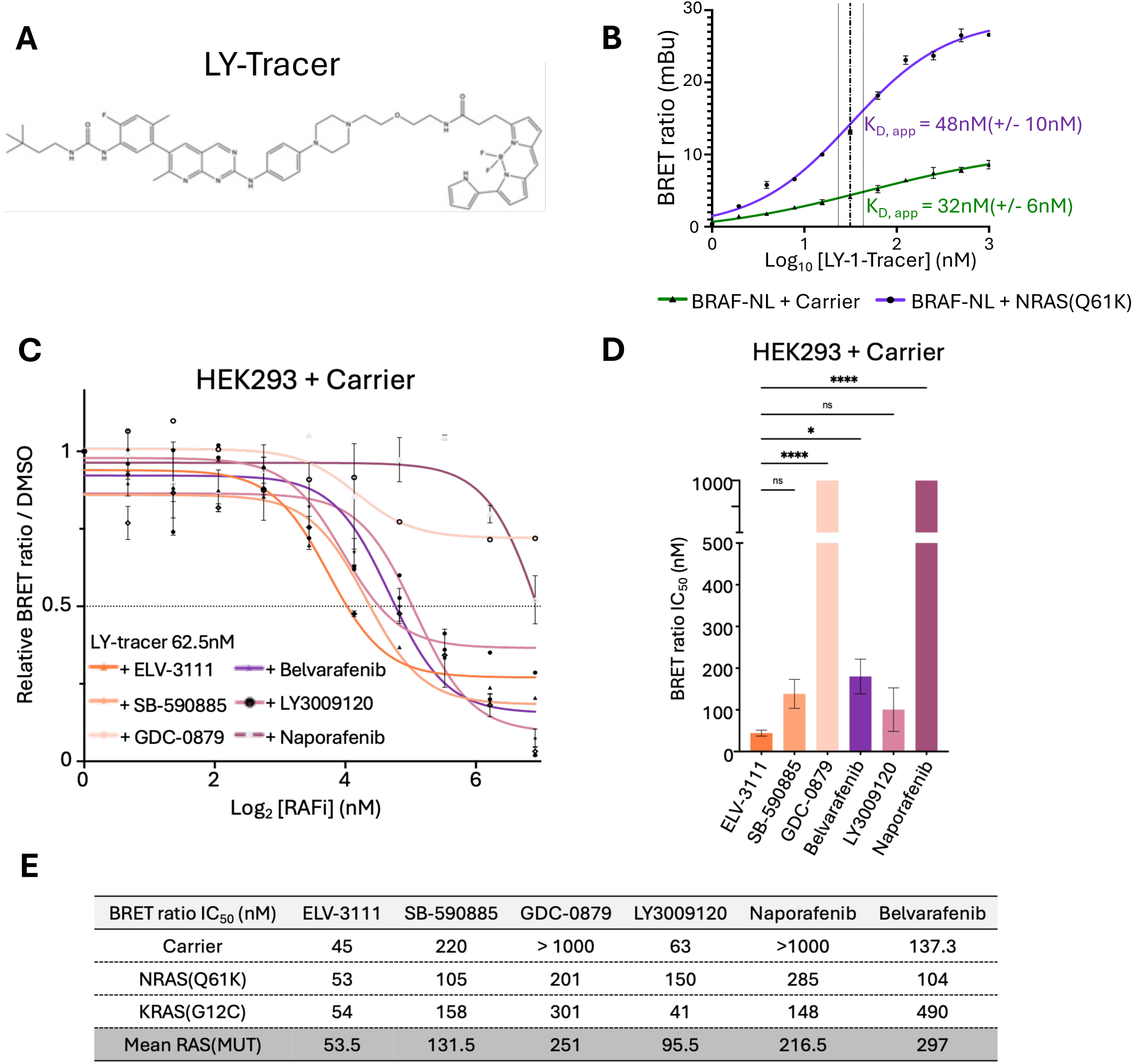
ELV-3111 is a Type 1 RAF inhibitor with broad activity and potent inhibition of both monomeric and dimeric RAF. (**A**) Structure of LY-Tracer used in NanoBRET experiments, derived from the Type 2 RAFi LY3009120. (**B**) HEK293 cells were co-transfected with BRAF-NanoLuc with or without NRAS(Q61K) for 24 hours. The following day, cells were treated with increasing concentration of LY-Tracer for 2 hours, followed by incubation with 20μM Fluorofurimazine, and BRET signal was measured. Data are presented as BRET ratio (*n=3* replicates). (**C**) HEK293 cells were co-transfected with BRAF-NanoLuc and plasmid-carrier and then treated with 62.5nM of LY-Tracer and increasing concentrations of different types of RAFis for 2 hours. After treatment, cells were incubated with 20μM Fluorofurimazine, and BRET signal was measured. Data are presented as relative BRET ratio normalized to DSMO-treated controls (mean values ± SEM, *n* = 3 independent experiments). (**D**) BRET ratio IC_50s_ values were calculated and represented as bar graphs (mean values ± SEM, *n* = 3 replicates. ns = not significant (P>0.05), * P<0.05, ****P<0.001, ordinary one-way ANOVA) or (**E)** as a table.

**Fig. S6.**
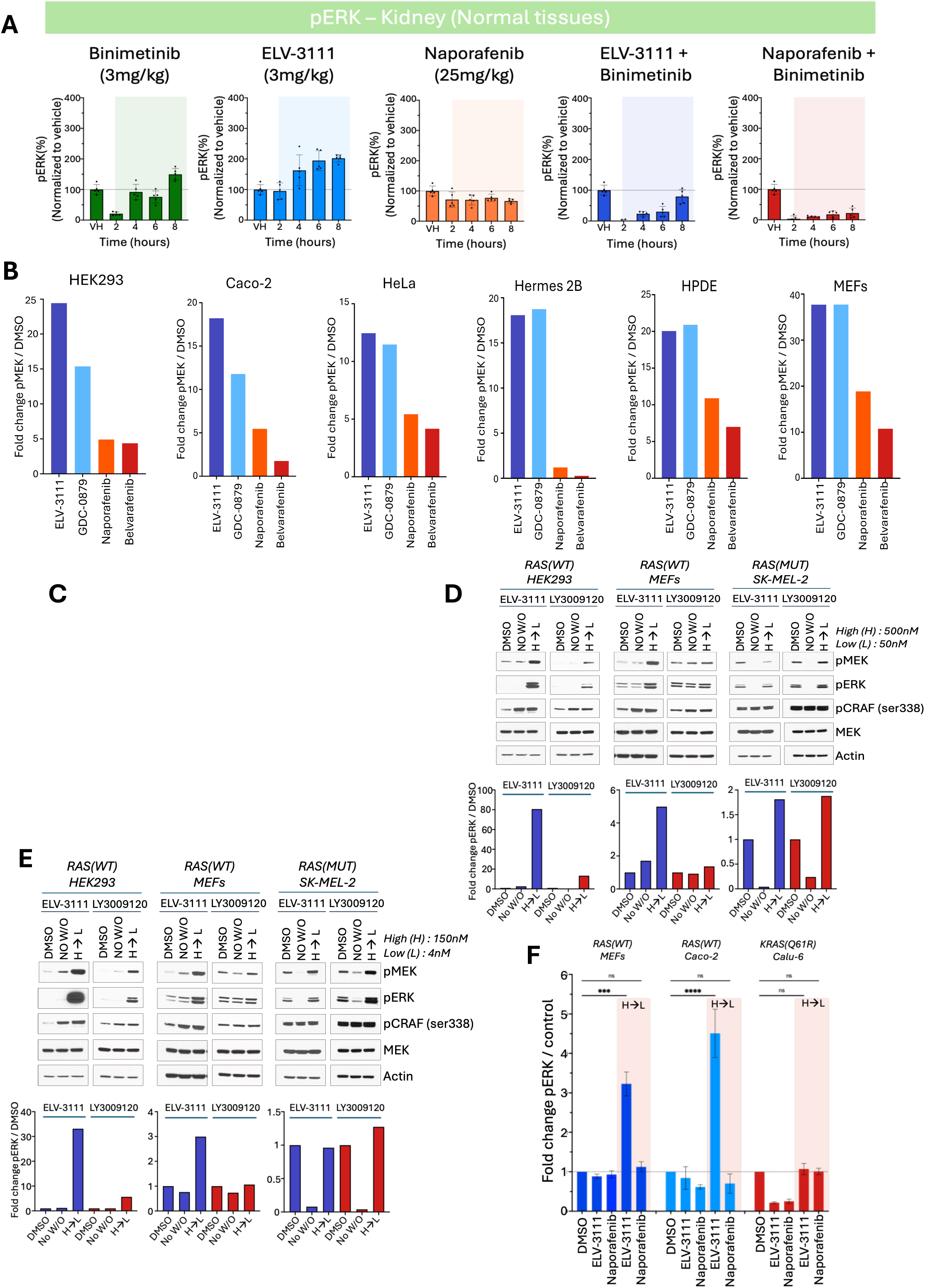
ELV-3111 potently suppresses MAPK in BRAF(MUT) and RAS(MUT) tumor models while inducing MAPK hyperactivation in normal tissues, *in vivo*. (**A**) Pharmacodynamic effect of ELV-3111 in mouse kidneys. SK-MEL-30 tumor xenograft-bearing mice were treated p.o with a single dose of 3mg/kg Binimetinib (MEKi), 3mg/kg ELV-3111 (Type 1 RAFi), 25mg/kg Naporafenib (Type 2 RAFi), or combinations of Binimetinib + ELV-3111 or Binimetinib + Naporafenib. Kidneys were harvested after 2, 4, 6 or 8 hours, lysed, and analyzed by immunoblotting for phosphorylated ERK (pERK) to assess MAPK pathway activity. Data represent mean relative pERK levels normalized to vehicle-treated controls (*n* = 5 mice per group). (**B**) A panel of RAS(WT) cell lines (HEK293, Caco-2, HeLa, Hermes 2B, HPDE, MEFs) was treated with 1μM of either Type 1 RAFi ELV-3111, Type 2 RAFis (Naporafenib or Belvarafenib), or 10μM of the Type 1 RAFi GDC-0879 for 8 hours. Following treatment, cells were washed and incubated in drug-free conditioned medium for 15 or 30 minutes, depending on the cell lines. Cell lysates were analyzed by immunoblotting to assess MAPK signaling. Fold change of phosphorylated MEK to MEK was calculated, compared to DMSO-treated controls, and represented as bar graphs. (**C**) Chemical structure of ELV-1974. (**D**) RAS(WT) HEK293 or MEFs cells and NRAS(Q61R) SK-MEL-2 were treated with 500nM (High – H) of Type 1 RAFi ELV-3111 or Type 2 RAFi LY3009120. Following treatment, cells were washed and incubated with 50nM (Low – L) of Type 1 RAFi ELV-3111 or Type 2 RAFi LY3009120. Cell lysates were analyzed by immunoblotting (**top**) with the indicated antibodies to assess MAPK signaling. MEK and actin were used as loading controls. Ratios of phosphorylated ERK to MEK for each sample were calculated and represented as fold change pERK bar graphs compared to DMSO controls (**bottom**). (**E**) RAS(WT) HEK293 or MEFs and NRAS(Q61R) SK-MEL-2 cells were treated with 150nM (High - H) of Type 1 ELV-3111 or Type 2 LY3009120. Following treatment, cells were washed and incubated with 4nM (Low - L) of Type 1 ELV-3111 or Type 2 LY3009120. Cell lysates were immunoblotted with the indicated antibodies to assess MAPK signaling. MEK and actin were used as loading controls. Ratios of phosphorylated ERK to MEK were calculated and represented as bar graphs compared to DMSO-treated controls. (**F**) RAS(WT) MEFs or Caco-2 cells and NRAS(Q61R) SK-MEL-2 were treated with 500nM (High – H) of Type 1 RAFi ELV-3111 or Type 2 RAFi Naporafenib. Following treatment, cells were washed and incubated with 50nM (Low – L) of Type 1 RAFi ELV-3111 or Type 2 RAFi Naporafenib. MAPK signaling pathway activation was assessed using TR-FRET-based detection assay for phospho-ERK (pERK) (mean values ± SEM, *n* = 3 replicates. ns = not significant (P>0.05), ***P<0.001, ****P<0.0001, ordinary one-way ANOVA).

**Fig S7.**
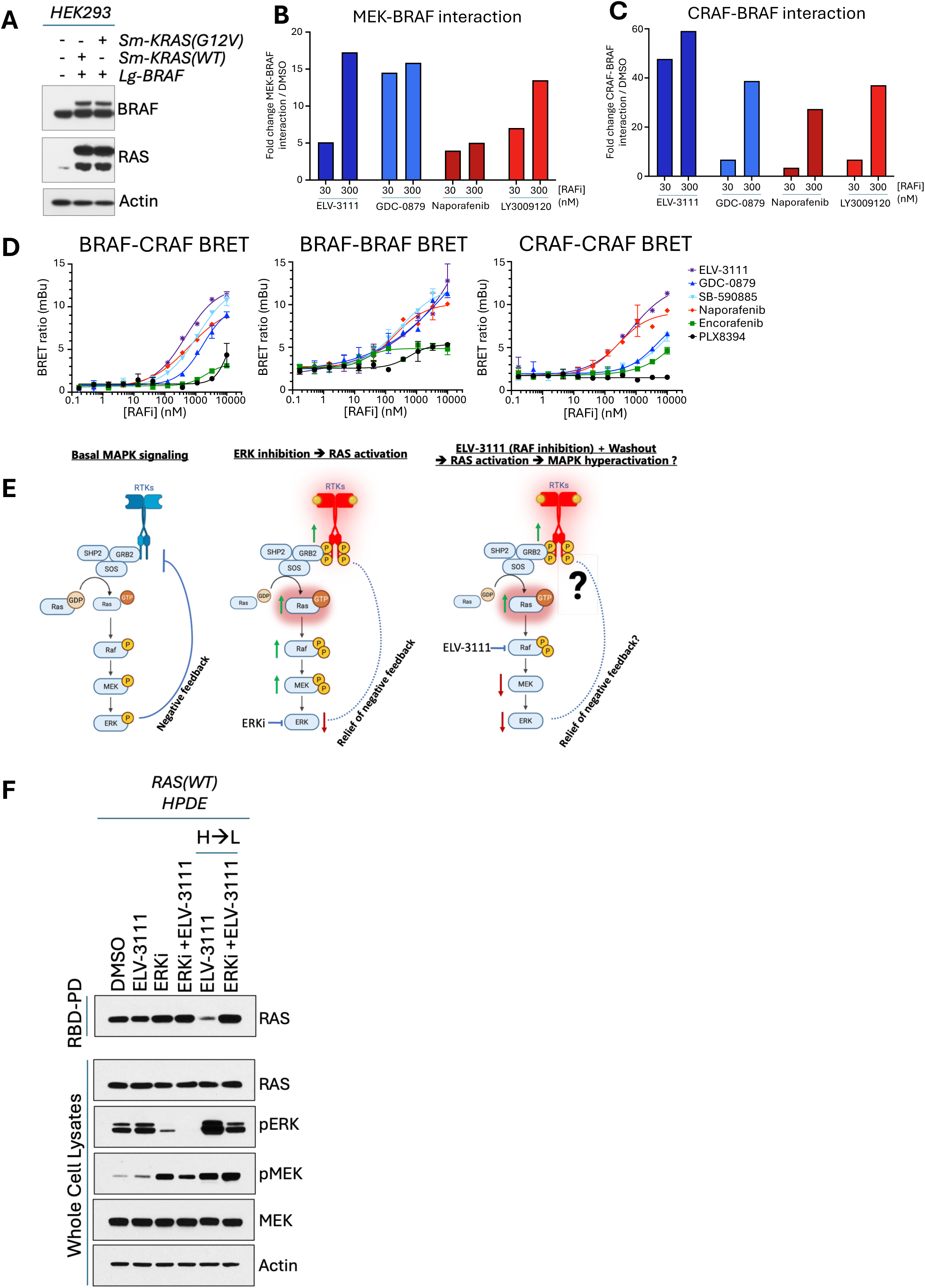
Type 1 RAFi-Induced MAPK hyperactivation is RAS-dependent. (**A**) HEK293 cells were co-transfected with LgBit-BRAF and either SmBit-KRAS(WT) or SmBit-KRAS(G12V) for 24 hours. Cell lysates were immunoblotted with the indicated antibodies to assess protein expression. Actin was used as loading control. (**B**) NRAS(Q61R) SK-MEL-2 cells were treated with different concentrations (0.03, 0.3μM) of the indicated RAFis for 1 hour. BRAF was immunoprecipitated, immunoblotted with the indicated antibodies to assess BRAF-CRAF dimerization and BRAF-MEK interaction, and quantified. Fold change (**B**) MEK-BRAF and (**C**) CRAF-BRAF interaction compared to DMSO-treated control were represented as bar graphs. (**D**) HEK293 cells transiently expressing a Nano-Luc and Halo-tagged pair of RAF proteins (BRAF:NanoLuc and CRAF:HaloTag for heterodimer assay) were treated with the 618 ligand and a concentration range of the indicated inhibitors for 2 hours. BRET ratio were measured and represented as curves (mean values± SEM, *n* = 3 independent experiments). (**E**) Schematic representation of the experimental hypothesis. (**F**) RAS(WT) HPDE cells were treated with 40nM (High - H) of Type 1 ELV-3111, 2μM of ERKi (Temuterkib), or the combination. Following treatment, cells were washed and incubated with 4nM (Low - L) of ELV-3111 in combination or not with 2μM of ERKi (Temuterkib). Cell lysates were subjected to RAS pull-down assay to assess RAS activation. Whole cell lysates were immunoblotted with the indicated antibodies to assess MAPK signaling. MEK and actin were used as loading controls.

**Fig. S8.** Type 1 RAFis exhibit over an order-of-magnitude faster dissociation from RAF relative to Type 2 RAFis. (**A**) The binding kinetics of Type 1 RAF inhibitors (ELV-3111, GDC-0879) and Type 2 RAF inhibitors (Naporafenib, LY3009120) were characterized by determining their association rate constant (k₁), dissociation rate constant (k₂), residence time (τ), and equilibrium dissociation constant (Kd). (**B**) Residence time (τ) for each RAFis were represented as bar graphs. (**C**) Corresponding *in-cell* residence time experiment, presented as bar graphs showing BRET ratios measured after 60 minutes (mean values ± SEM, *n* = 3 independent experiments. ****P<0.0001, ordinary one-way ANOVA). (**D and E**) Molecular dynamics (MD) simulations of the Type 2 RAFi Sorafenib and the Type 1 RAFi GDC-0879. Shown are ligand RMSD over 1 μs, ligand chemical structures with corresponding ligand RMSF profiles, superposition of ligand conformations sampled at 200, 400, 600, and 800 ns from the 1 μs trajectories, illustrating stable binding within the BRAF active site relative to the hinge region, αC-helix, and activation loop, and protein–ligand contact frequency plots summarizing hydrogen bonds, hydrophobic interactions, ionic interactions, and water bridges observed throughout the simulations. (**F**) Two-dimensional ligand interaction diagrams summarizing the key residues mediating contacts with Type 2 RAFis (LY3009120 and Sorafenib) or Type 1 RAFis (ELV-3111 and GDC-0879).

**Fig. S9.**
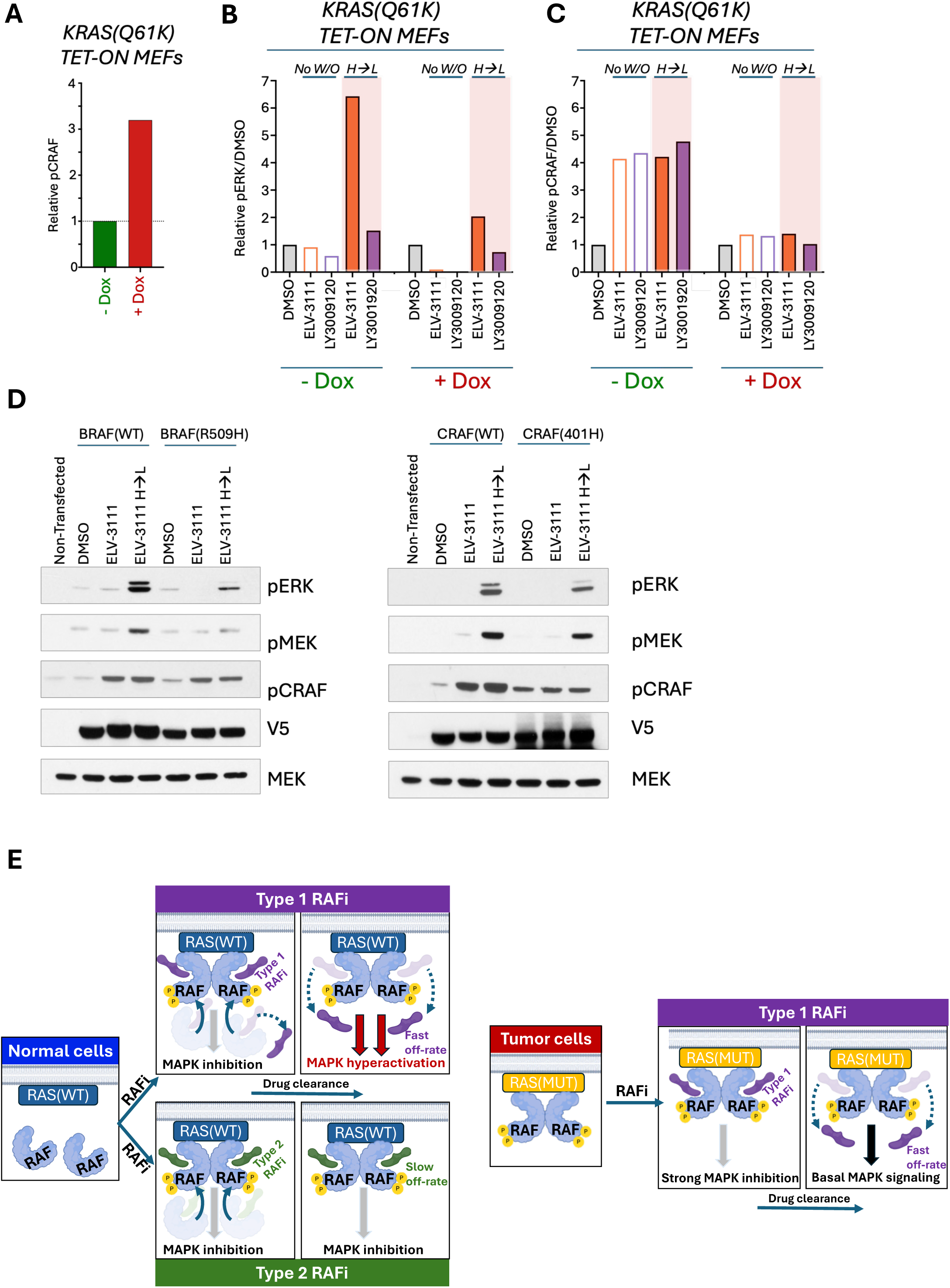
Type 1 RAFis exhibit over an order-of-magnitude faster dissociation from RAF relative to Type 2 RAFis. MEFs cells engineered to express doxycycline-inducible KRAS(Q61K) were treated with doxycycline (1μg/mL) for 48 hours, followed by treatment with 500nM (High – H) of Type 1 RAFi (ELV-3111) or Type 2 RAFi (LY3009120) for 8 hours. Following treatment, cells were washed and treated with 50nM (Low – L) of either ELV-3111 or LY3009120. Cell lysates were analyzed by immunoblotting, similarly to Fig. 4G. (**A**) pCRAF levels at baseline and after or without doxycycline induction were quantified and represented as a bar graph. (**B**) pERK and (**C**) pCRAF levels following RAFi treatments were quantified from the immunoblots, normalized to DMSO control, and represented as bar graphs. (**D**) HEK293 cells ectopically expressing V5-BRAF(WT), V5-BRAF(509H), V5-CRAF(WT), or V5-CRAF(401H) were treated with 500nM (High – H) of Type 1 RAFi ELV-3111 for 8 hours. Following treatment, cells were washed and treated with 50nM (Low – L) of ELV-3111. Cell lysates were analyzed by immunoblotting to assess MAPK signaling. MEK was used as loading control. (**E**) Model of MAPK hyperactivation mechanism mediated by Type 1 RAFis in normal cells. Both Type 1 and Type 2 RAFis induce RAF-active conformation and promote RAF “priming” upon binding. However, due to rapid dissociation (short residence time), Type 1 RAFis induce MAPK hyperactivation upon drug withdrawal in normal cells. In contrast, Type 2 RAFis, which have longer residence time, do not elicit this hyperactivation. In RAS(MUT) setting, RAF is constitutively in active conformation and treatment with Type 1 RAFis does not induce MAPK hyperactivation.

**Fig. S10.**
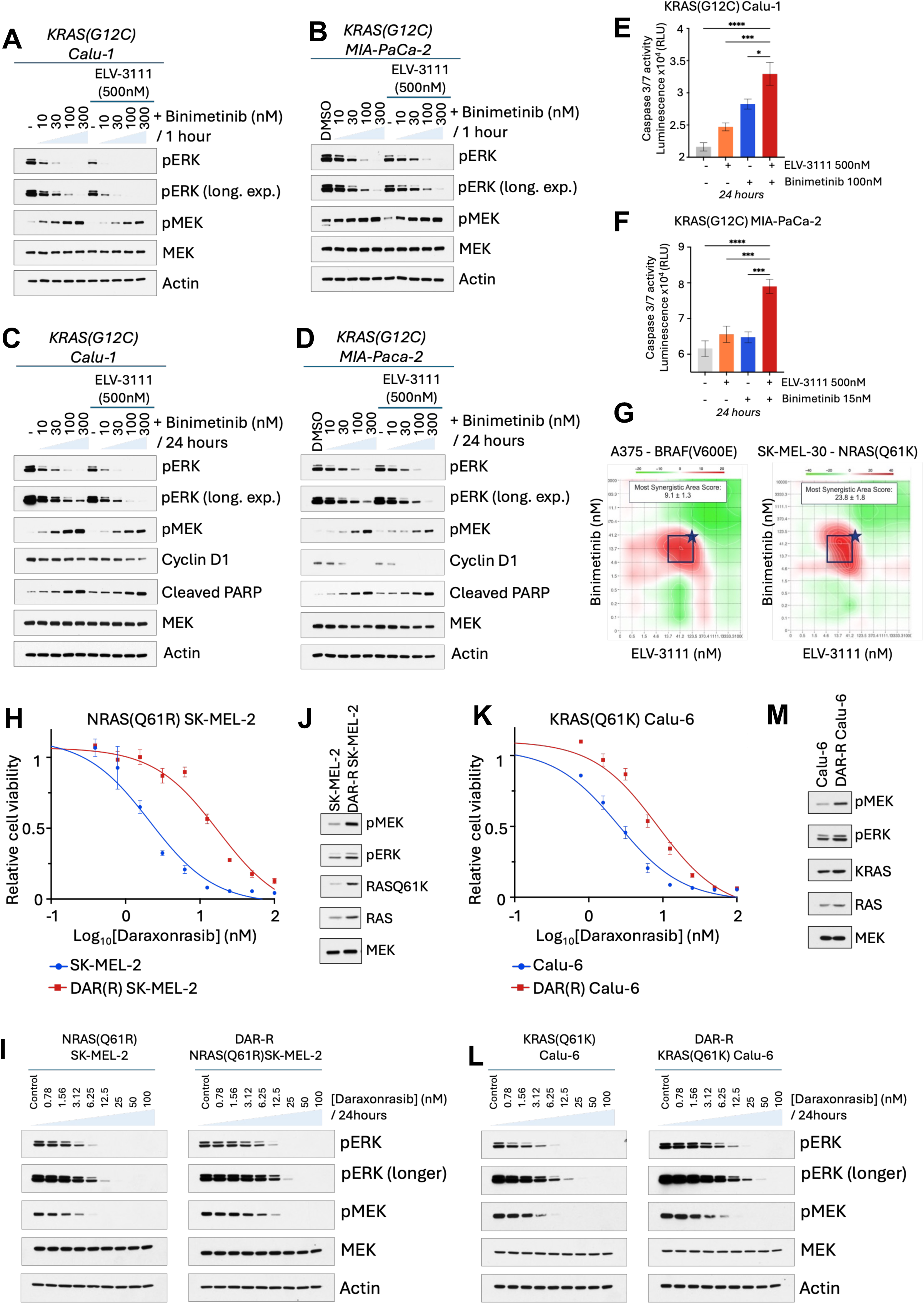
ELV-3111 – based MAPK targeting combinations are highly potent in BRAF(MUT) and RAS(MUT) tumors and well tolerated *in vivo*. KRAS(G12C) (**A**) Calu-1 or (**B**) MIA-PaCa-2 were treated for 1 hour with increasing concentration of Binimetinib (MEKi) in combination or not with 500nM of Type 1 RAFi ELV-3111. Cell lysates were analyzed by immunoblotting with the indicated antibodies to assess MAPK signaling. MEK and actin were used as loading controls. KRAS(G12C) (**C**) Calu-1 or (**D**) MIA-PaCa-2 were treated for 24 hours with increasing concentrations of Binimetinib (MEKi) in combination or not with 500nM of Type 1 RAFi ELV-3111. Cell lysates were analyzed by immunoblotting with the indicated antibodies to assess MAPK signaling. MEK and actin were used as loading controls. KRAS(G12C) (**E**) Calu-1 or (**F**) MIA-PaCa-2 were treated with 100nM or 15nM of Binimetinib (MEKi), 500nM of Type 1 RAFi ELV-311, or the combination. Apoptosis of cells was assessed measuring Caspase 3/7 activity after 24 hours of treatment (mean values ± SEM, *n* = 3 replicates, <u>*P<0.05, ***P<0.001, ****P<0.0001, ordinary one-way ANOVA</u>). (**G**) A375 or SK-MEL-30 cells were treated with a combination dose range of ELV-3111 and Binimetinib for 72h (A375) or 120h (SK-MEL-30). Relative cell viability was measured, uploaded into SynergyFinder 3.0 and scored using the highest single agent (HSA) synergistic area method. Star indicates intersection of single agent GI_50_s. (**H**) Parental NRAS(Q61R) SK-MEL-2 or Daraxonrasib-resistant NRAS(Q61R) SK-MEL-2 (DAR-R SK-MEL-2) cells were treated for 6 days with increasing concentrations of Daraxonrasib (RASONi). Cell viability was measured and represented as relative viability compared to DMSO-treated controls (mean values ± SEM, *n* = 3 independent experiments). (**I**) Parental NRAS(Q61R) SK-MEL-2 or Daraxonrasib-resistant NRAS(Q61R) SK-MEL-2 (DAR-R SK-MEL-2) cells were treated for 24 hours with increasing concentrations of Daraxonrasib (RASONi). Cell lysates were analyzed by immunoblotting with the indicated antibodies to assess MAPK signaling. MEK and actin were used as loading controls. (**J**) Parental NRAS(Q61R) SK-MEL-2 or Daraxonrasib-resistant NRAS(Q61R) SK-MEL-2 (DAR-R SK-MEL-2) cells were analyzed by immunoblotting with the indicated antibodies to assess RAS level expression and MAPK signaling. MEK was used as loading control. (**K**) Parental KRAS(Q61K) Calu-6 or Daraxonrasib-resistant KRAS(Q61K) Calu-6 (DAR-R Calu-6) cells were treated for 6 days with increasing concentrations of Daraxonrasib (RASONi). Cell viability was measured and represented as relative viability compared to DMSO-treated controls (mean values ± SEM, *n* = 3 independent experiments). (**L**) Parental KRAS(Q61K) Calu-6 or Daraxonrasib-resistant KRAS(Q61K) Calu-6 (DAR-R Calu-6) cells were treated for 24 hours with increasing concentrations of Daraxonrasib (RASONi). Cell lysates were analyzed by immunoblotting with the indicated antibodies to assess MAPK signaling. MEK and actin were used as loading controls. (**M**) Parental KRAS(Q61K) Calu-6 or Daraxonrasib-resistant KRAS(Q61K) Calu-6 (DAR-R Calu-6) cells were analyzed by immunoblotting with the indicated antibodies to assess RAS level expression and MAPK signaling. MEK was used as loading control.

**Fig. S11.**
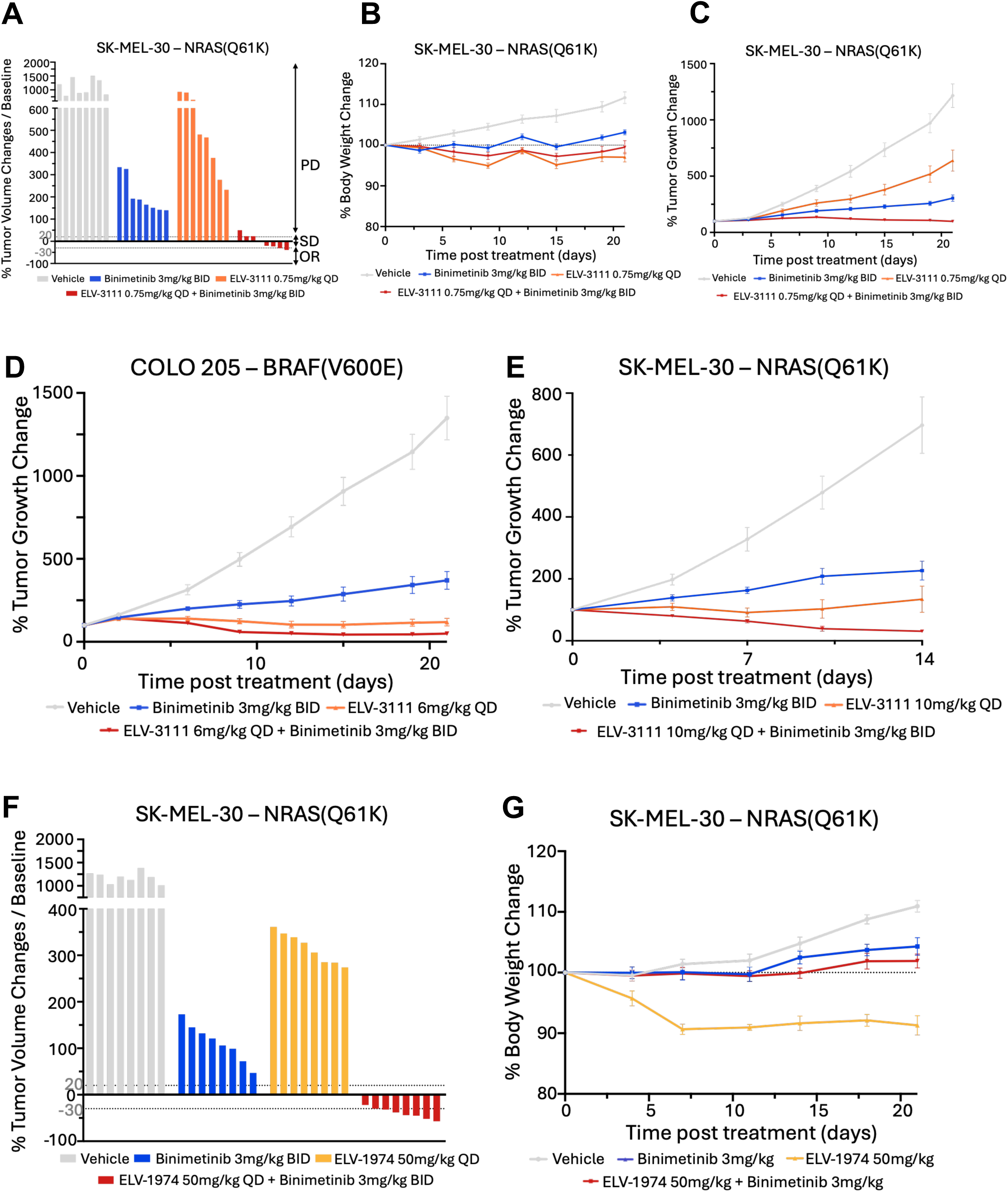
ELV-3111 – based MAPK targeting combinations are highly potent in a tumor model insensitive to current MAPK-targeting therapies. Mice with established SK-MEL-30 (NRAS(Q61K)) tumors were treated p.o. with vehicle, 3mg/kg Binimetinib BID, 0.75mg/kg ELV-3111 QD or the combination (*n* =8 mice per group). (**A**) Percent change in individual tumor volume at the end of study, (**B**) body weight of mice and (**C**) tumor growth changes were followed over time and normalized to the baseline. (**D**) and (**E**) Mice with established COLO 205 (BRAF(V600E)) or SK-MEL-30 (NRAS(Q61K)) tumors were treated p.o. with vehicle, 3 mg/kg Binimetinib BID, 6 mg/kg (COLO-205) or 10 mg/kg (SK-MEL-30) ELV-3111 QD or the combination, when tumors reached ∼150mm^3^. Tumor growth changes over time were followed and normalized to the baseline. Mice with established SK-MEL-30 (NRAS(Q61K)) tumors were treated p.o. with vehicle, 3 mg/kg Binimetinib BID, 50mg/kg ELV-1974 QD or the combination (*n* = 8 mice/group). (**F**) Percent change in individual tumor volume at the end of the study, (**G**) body weight of mice over time were followed and normalized to the baseline.

**Fig. S12.**
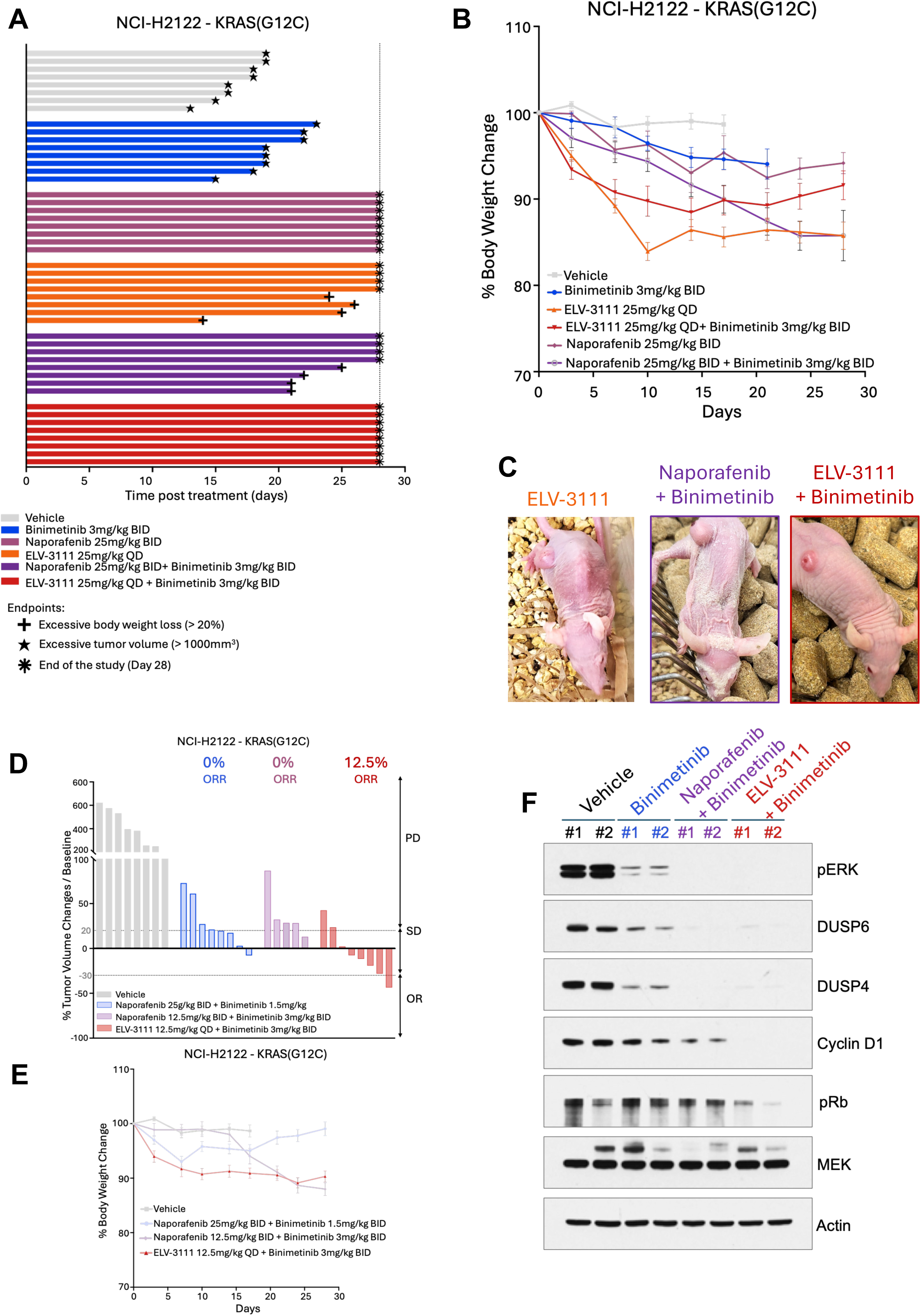
ELV-3111 – based MAPK targeting combinations are highly potent in a tumor model insensitive to current MAPK-targeting therapies. Mice with established NCI-H2122 (KRAS(G12C)) tumors (∼250 mm³) were treated orally with vehicle, Binimetinib (3mg/kg BID), Naporafenib (25mg/kg BID), ELV-3111 (25mg/kg QD), or the combinations of Naporafenib + Binimetinib and ELV-3111 + Binimetinib (*n = 8* mice per group). (**A**) Swimmer plot showing individual treatment durations and event timelines for each animal and (**B**) body weight of mice over time was followed overtime and normalized to the baseline. (**C**) Representative images of mice under different treatment conditions. Mice with established NCI-H2122 (KRAS(G12C)) tumors (∼250mm3), were treated p.o. with vehicle, 1.5mg/kg Binimetinib BID in combination with 25mg/kg Naporafenib BID, 3mg/kg Binimetinib BID in combination with 12.5mg/kg Naporafenib BID or 3mg/kg Binimetinib BID in combination with 12.5mg/kg ELV-3111 QD (*n = 8* mice per group). (**D**) Percent change in individual tumor volume at the end of study and (**E**) body weight of mice over time were followed and normalized to the baseline. (**F**) Mice with established NCI-H2122 (KRAS(G12C)) tumors (∼500mm3), were treated p.o. for 72 hours with vehicle, 3mg/kg Binimetinib BID, in combination or not with Naporafenib (25mg/kg BID), or ELV-3111 (25mg/kg QD) (*n* = 2 mice/group). Tumors were harvested after 72 hours of treatment, lysed, and analyzed by immunoblotting to assess MAPK pathway activity and downstream signaling. MEK and actin were used as loading controls.

**Table S1.** Crystallographic data collection and refinement statistics.

|  | **BRAF kinase domain**<br>**bound to ELV-3111** |
| --- | --- |
| PDB ID | 9ZBO |
| **Data collection** |  |
| Space group | P 4<sub>1</sub> 2<sub>1</sub> 2 |
| Cell dimensions |  |
| a, b, c (Å) | 94.24, 94.24, 162.74 |
| α, β, γ (°) | 90.0, 90.0, 90.0 |
| Resolution (Å) | 61.74–2.85 (2.90–2.85)* |
| Rsym | 0.190 (2.490) |
| Rpim | 0.052 (0.657) |
| Rmeas | 0.198 (2.576) |
| Mean I/σ(I) | 11.5 (1.3) |
| Completeness (%) | 98.8 (100.0) |
| Multiplicity | 14.4 (15.1) |
| CC1/2 (%) | 99.8 (55.0) |
| **Refinement** |  |
| Resolution (Å) | 61.74 - 2.85 |
| No. Reflections (working/test) | 16953/700 |
| Rcryst / Rfree (%) | 21.7/26.4 |
| No. atoms |  |
| Protein | 4323 |
| Ligand/ion | 80 |
| Water | 5 |
| B factors (Å<sup>2</sup>) |  |
| Protein | 80.2 |
| Ligand/ions | 65.3 |
| Water | 56.8 |
| Other atoms | 102.2 |
| R.M.S. deviations |  |
| Bond length (Å) | 0.003 |
| Bond angles (°) | 1.32 |
| Ramachandran plot |  |
| Favored (%) | 96.22 |
| Allowed (%) | 3.59 |
| Outliers (%) | 0.19 |

\* Values in parentheses are for the highest-resolution shell.

**Table S2.**
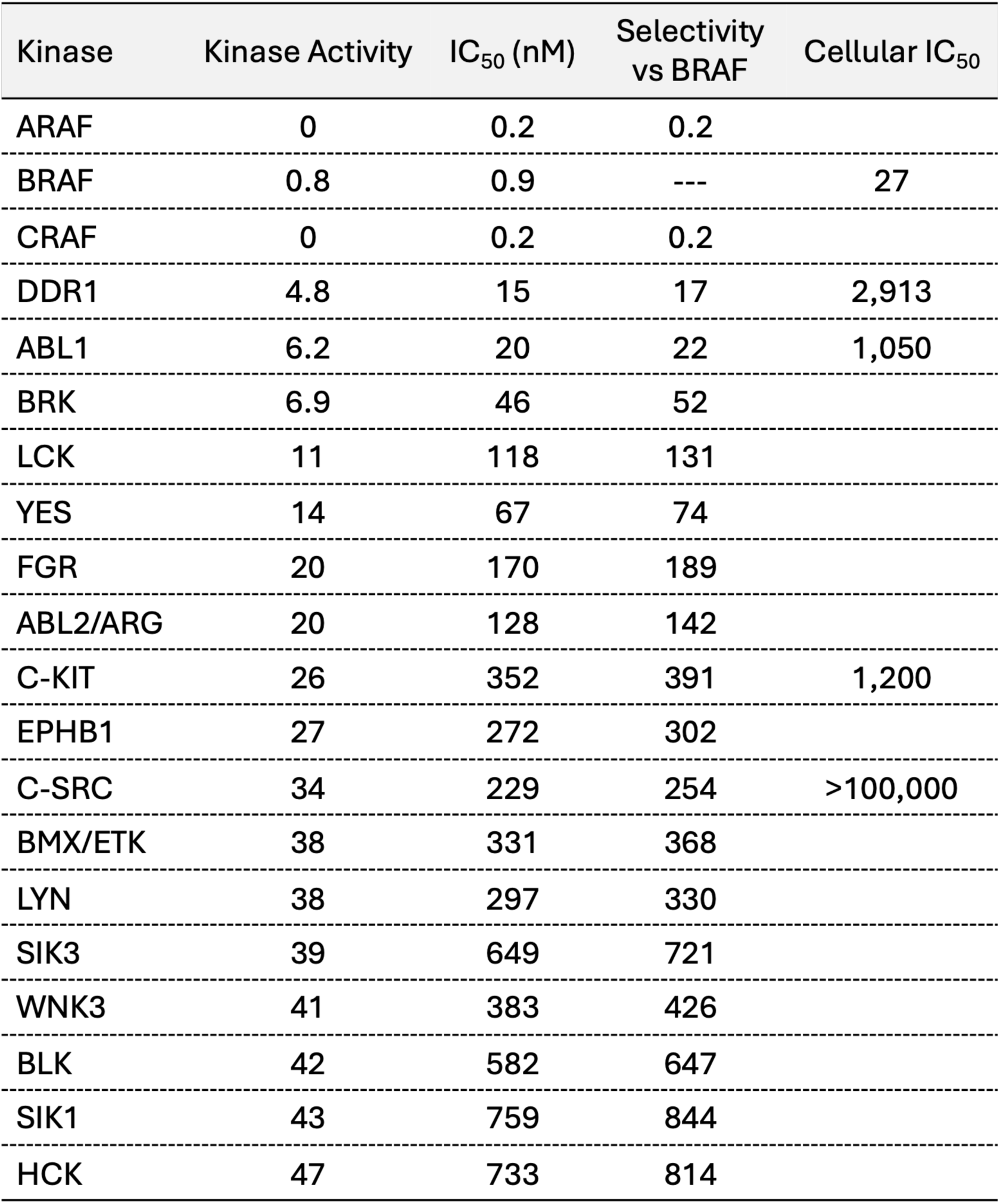
Kinome selectivity of ELV-3111. Kinases identified from the screen as being inhibited more than 50% with 1 μM ELV-3111 were selected for follow-up IC_50_ determinations with a dose range of ELV-3111. Selectivity was determined by comparing anti-target IC_50_s to the IC_50_s for BRAF. For select kinases with known potential toxicities when _inhibited_ in the clinic, we also calculated cellular IC_50_s. For BRAF pERK inhibition was measured by in-cell western, for DDR1 target engagement was measured by NanoBRET, for ABL1 inhibition of the K562 cell line proliferation was measured with Cell-Titer Glo, for C-KIT and C-SRC inhibition of autophosphorylation was measured by ELISA.

**Table S3.** Comparing the activity of ELV-3111 to other MAPK pathway inhibitors in BRAF and NRAS mutant cell lines. Cell lines with a spectrum of BRAF and NRAS mutations were screened for sensitivity to ELV-3111 and other MAPK pathway inhibitors. An in-cell western procedure was used to measure pERK inhibition after 2 hours of treatment with a dose range of compounds. Proliferation assays were performed in the indicated cell lines after 72h (A375) or 120h (all other lines). pERK IC_50_s or GI_50_s are represented as table (geometric mean, *n* = 3 independent experiments).

|  |  | Class I | Class II |  |  | Class III |  | NRAS Melanoma |  |  |  |
| --- | --- | --- | --- | --- | --- | --- | --- | --- | --- | --- | --- |
| pERK IC<sub>50</sub> (nM) |  | A375<br>V600E | WM1789<br>K601E | WM3928<br>SKAP2-BRAF | WM3670<br>G469E/<br>NRAS G12D | WM3912<br>N581Y | WM3629<br>D594G/<br>NRAS G12D | HMVII<br>Q61L (BRAF G469V) | SK-MEL-30<br>Q61K | WM3406<br>Q61K | WM3854<br>Q61L |
| MEKi | Binimetinib | 17 | 38 | 7 | 21 | 19 | 17 | 16 | 7 | 4 | 6 |
| Type 1.5 | Dabrafenib | 3 | 32 | 1,110 | 7,890 | 3,850 | 2,580 | 1,770 | 3,180 | 2,300 | >5,000 |
|  | Encorafenib | 6 | 26 | 360 | >3,000 | 1,140 | 1,540 | 694 | 923 | 1,120 | 1,800 |
| Type 2 | Naporafenib | 416 | 216 | 179 | 20 | 31 | 9 | 8 | 38 | 57 | 96 |
|  | Belvarafenib | 87 | 381 | 328 | 130 | 195 | 59 | 39 | 42 | 88 | 170 |
| Type 1 | ELV-3111 | 21 | 93 | 122 | 19 | 24 | 2 | 24 | 30 | 29 | 60 |
| GI<sub>50</sub> (nM) |  | A375<br>V600E | WM1789<br>K601E | WM3928<br>SKAP2-BRAF | WM3670<br>G469E/<br>NRAS G12D | WM3912<br>N581Y | WM3629<br>D594G/<br>NRAS G12D | HMVII<br>Q61L (BRAF G469V) | SK-MEL-30<br>Q61K | WM3406<br>Q61K | WM3854<br>Q61L |
| MEKi | Binimetinib | 16 | 162 | 14 | 3630 | 5070 | 335 | 43 | 52 | 15 | 18 |
| Type 1.5 | Dabrafenib | 3 | 46 | 1,250 | >25,000 | >25,000 | 2,180 | >1,560 | >10,000 | >7,190 | >10,000 |
|  | Encorafenib | 3 | 11 | 632 | >5,000 | 2,320 | 1,520 | 563 | 1,690 | 614 | 834 |
| Type 2 | Naporafenib | 245 | 222 | 95 | 1,250 | 1,920 | 47 | 12 | 123 | 86 | 75 |
|  | Belvarafenib | 67 | 106 | 121 | >25,000 | 8,230 | 109 | 29 | 125 | 41 | 35 |
| Type 1 | ELV-3111 | 42 | 1460 | 239 | 950 | 526 | 73 | 34 | 125 | 109 | 125 |

## References in Materials and Methods

1. P. Emsley, K. Cowtan, Coot: Model-building tools for molecular graphics. Acta Crystallogr. D Biol. Crystallogr. 60, 2126–2132 (2004).

2. Winn, M. D. et al. “Overview of the CCP4 suite and current developments.” *Acta Cryst. D67*, 235-242 (2011)

3. Vonrhein, C., Flensburg, C., Keller, P., Sharff, A., Smart, O., Paciorek, W., Womack, T. & Bricogne, G. (2011). Data processing and analysis with the autoPROC toolbox. Acta Crystallographica Section D: Structural Biology, 67(4), 293–302.

4. Ianevski, A., Giri, K. A., Aittokallio, T., 2022. SynergyFinder 3.0: an interactive analysis and consensus interpretation of multi-drug synergies across multiple samples. Nucleic Acids Research. gkac382, 10.1093/nar/gkac382

## References

1. A. Young et al., Ras signaling and therapies. Advances in cancer research 102, 1–17 (2009).

2. A. R. Moore, S. C. Rosenberg, F. McCormick, S. Malek, RAS-targeted therapies. Nature reviews. Drug discovery, (2021).

3. A. A. Samatar, P. I. Poulikakos, Targeting RAS-ERK signalling in cancer: promises and challenges. Nature reviews. Drug discovery 13, 928–942 (2014).

4. J. Zhang, P. L. Yang, N. S. Gray, Targeting cancer with small molecule kinase inhibitors. Nature reviews. Cancer 9, 28–39 (2009).

5. Z. Karoulia et al., An Integrated Model of RAF Inhibitor Action Predicts Inhibitor Activity against Oncogenic BRAF Signaling. Cancer cell 30, 485–498 (2016).

6. Z. Karoulia, E. Gavathiotis, P. I. Poulikakos, New perspectives for targeting RAF kinase in human cancer. Nature reviews. Cancer 17, 676–691 (2017).

7. J. Tsai et al., Discovery of a selective inhibitor of oncogenic B-Raf kinase with potent antimelanoma activity. Proceedings of the National Academy of Sciences of the United States of America 105, 3041–3046 (2008).

8. P. I. Poulikakos, C. Zhang, G. Bollag, K. M. Shokat, N. Rosen, RAF inhibitors transactivate RAF dimers and ERK signalling in cells with wild-type BRAF. Nature 464, 427–430 (2010).

9. E. W. Joseph et al., The RAF inhibitor PLX4032 inhibits ERK signaling and tumor cell proliferation in a V600E BRAF-selective manner. Proceedings of the National Academy of Sciences of the United States of America 107, 14903–14908 (2010).

10. G. Hatzivassiliou et al., RAF inhibitors prime wild-type RAF to activate the MAPK pathway and enhance growth. Nature 464, 431–435 (2010).

11. K. T. Flaherty et al., Combined BRAF and MEK inhibition in melanoma with BRAF V600 mutations. The New England journal of medicine 367, 1694–1703 (2012).

12. C. Robert et al., Improved overall survival in melanoma with combined dabrafenib and trametinib. The New England journal of medicine 372, 30–39 (2015).

13. R. Dummer et al., Encorafenib plus binimetinib versus vemurafenib or encorafenib in patients with BRAF-mutant melanoma (COLUMBUS): a multicentre, open-label, randomised phase 3 trial. The lancet oncology 19, 603–615 (2018).

14. S. Kopetz et al., Encorafenib, Binimetinib, and Cetuximab in BRAF V600E-Mutated Colorectal Cancer. The New England journal of medicine 381, 1632–1643 (2019).

15. I. Yen et al., ARAF mutations confer resistance to the RAF inhibitor belvarafenib in melanoma. Nature 594, 418–423 (2021).

16. R. J. Sullivan et al., A Phase I Study of LY3009120, a Pan-RAF Inhibitor, in Patients with Advanced or Metastatic Cancer. Molecular cancer therapeutics 19, 460–467 (2020).

17. F. de Braud et al., Initial Evidence for the Efficacy of Naporafenib in Combination With Trametinib in NRAS-Mutant Melanoma: Results From the Expansion Arm of a Phase Ib, Open-Label Study. Journal of clinical oncology : official journal of the American Society of Clinical Oncology 41, 2651–2660 (2023).

18. M. B. Robers et al., Target engagement and drug residence time can be observed in living cells with BRET. Nature communications 6, 10091 (2015).

19. K. A. Monaco et al., LXH254, a Potent and Selective ARAF-Sparing Inhibitor of BRAF and CRAF for the Treatment of MAPK-Driven Tumors. Clinical cancer research : an official journal of the American Association for Cancer Research 27, 2061–2073 (2021).

20. K. P. Hoeflich et al., Antitumor efficacy of the novel RAF inhibitor GDC-0879 is predicted by BRAFV600E mutational status and sustained extracellular signal-regulated kinase/mitogen-activated protein kinase pathway suppression. Cancer research 69, 3042–3051 (2009).

21. C. Adamopoulos et al., Exploiting Allosteric Properties of RAF and MEK Inhibitors to Target Therapy-Resistant Tumors Driven by Oncogenic BRAF Signaling. Cancer discovery 11, 1716–1735 (2021).

22. P. I. Poulikakos et al., RAF inhibitor resistance is mediated by dimerization of aberrantly spliced BRAF(V600E). Nature 480, 387–390 (2011).

23. S. B. Peng et al., Inhibition of RAF Isoforms and Active Dimers by LY3009120 Leads to Anti-tumor Activities in RAS or BRAF Mutant Cancers. Cancer cell 28, 384–398 (2015).

24. S. Kopetz et al., Molecular profiling of BRAF-V600E-mutant metastatic colorectal cancer in the phase 3 BEACON CRC trial. Nature medicine 30, 3261–3271 (2024).

25. R. Nazarian et al., Melanomas acquire resistance to B-RAF(V600E) inhibition by RTK or N-RAS upregulation. Nature 468, 973–977 (2010).

26. J. D. Vasta et al., Quantitative, Wide-Spectrum Kinase Profiling in Live Cells for Assessing the Effect of Cellular ATP on Target Engagement. Cell Chem Biol 25, 206–214 e211 (2018).

27. T. A. Ahmed et al., SHP2 Drives Adaptive Resistance to ERK Signaling Inhibition in Molecularly Defined Subsets of ERK-Dependent Tumors. Cell Rep 26, 65–78 e65 (2019).

28. P. Lito et al., Relief of profound feedback inhibition of mitogenic signaling by RAF inhibitors attenuates their activity in BRAFV600E melanomas. Cancer cell 22, 668–682 (2012).

29. J. S. Duncan et al., Dynamic reprogramming of the kinome in response to targeted MEK inhibition in triple-negative breast cancer. Cell 149, 307–321 (2012).

30. R. B. Corcoran et al., EGFR-mediated re-activation of MAPK signaling contributes to insensitivity of BRAF mutant colorectal cancers to RAF inhibition with vemurafenib. Cancer discovery 2, 227–235 (2012).

31. C. Liu et al., Combinations with Allosteric SHP2 Inhibitor TNO155 to Block Receptor Tyrosine Kinase Signaling. Clinical cancer research : an official journal of the American Association for Cancer Research 27, 342–354 (2021).

32. A. Cuevas-Navarro et al., RAS-dependent RAF-MAPK hyperactivation by pathogenic RIT1 is a therapeutic target in Noonan syndrome-associated cardiac hypertrophy. Sci Adv 9, eadf4766 (2023).

33. A. S. Dixon et al., NanoLuc Complementation Reporter Optimized for Accurate Measurement of Protein Interactions in Cells. ACS Chem Biol 11, 400–408 (2016).

34. C. Zhang et al., RAF inhibitors that evade paradoxical MAPK pathway activation. Nature 526, 583–586 (2015).

35. T. Jin et al., RAF inhibitors promote RAS-RAF interaction by allosterically disrupting RAF autoinhibition. Nature communications 8, 1211 (2017).

36. D. S. Hong et al., KRAS(G12C) Inhibition with Sotorasib in Advanced Solid Tumors. The New England journal of medicine 383, 1207–1217 (2020).

37. P. A. Janne et al., Adagrasib in Non-Small-Cell Lung Cancer Harboring a KRAS(G12C) Mutation. The New England journal of medicine 387, 120–131 (2022).

38. R. Y. Ebright, J. Dilly, A. T. Shaw, A. J. Aguirre, Response and Resistance to RAS Inhibition in Cancer. Cancer discovery 15, 1325–1349 (2025).

39. M. Holderfield et al., Concurrent inhibition of oncogenic and wild-type RAS-GTP for cancer therapy. Nature, (2024).

40. J. Cregg et al., Discovery of Daraxonrasib (RMC-6236), a Potent and Orally Bioavailable RAS(ON) Multi-selective, Noncovalent Tri-complex Inhibitor for the Treatment of Patients with Multiple RAS-Addicted Cancers. Journal of medicinal chemistry 68, 6064–6083 (2025).

41. M. M. Awad et al., Acquired Resistance to KRAS(G12C) Inhibition in Cancer. The New England journal of medicine 384, 2382–2393 (2021).

42. R. Yaeger et al., Molecular Characterization of Acquired Resistance to KRASG12C-EGFR Inhibition in Colorectal Cancer. Cancer discovery 13, 41–55 (2023).

43. J. Dilly et al., Mechanisms of Resistance to Oncogenic KRAS Inhibition in Pancreatic Cancer. Cancer discovery 14, 2135–2161 (2024).

44. P. Cassier et al., 613MO A phase I clinical trial evaluating exarafenib, a selective pan-RAF inhibitor in combination with binimetinib in NRAS-mutant (NRASMut) melanoma (Mel) & BRAF-altered solid tumors. Annals of Oncology 35, S491–S492 (2024).

45. I. Yen et al., Pharmacological Induction of RAS-GTP Confers RAF Inhibitor Sensitivity in KRAS Mutant Tumors. Cancer cell 34, 611–625 e617 (2018).

46. M. H. Hofmann et al., BI-3406, a Potent and Selective SOS1-KRAS Interaction Inhibitor, Is Effective in KRAS-Driven Cancers through Combined MEK Inhibition. Cancer discovery 11, 142–157 (2021).

47. A. Tedeschi et al., Pan-KRAS Inhibitors BI-2493 and BI-2865 Display Potent Antitumor Activity in Tumors with KRAS Wild-type Allele Amplification. Molecular cancer therapeutics 24, 550–562 (2025).

48. T. Okuzumi et al., Inhibitor hijacking of Akt activation. Nature chemical biology 5, 484–493 (2009).

49. L. R. Pearce et al., Characterization of PF-4708671, a novel and highly specific inhibitor of p70 ribosomal S6 kinase (S6K1). Biochem J 431, 245–255 (2010).

